# Reverse chemical ecology approach for sustainable palm tree protection against invasive palm weevils

**DOI:** 10.1101/2023.01.13.523742

**Authors:** Binu Antony, Nicolas Montagné, Arthur Comte, Sara Mfarrej, Jernej Jakše, Rémi Capoduro, Rajan Shelke, Khasim Cali, Mohammed Ali AlSaleh, Krishna Persaud, Arnab Pain, Emmanuelle Jacquin-Joly

## Abstract

The reverse chemical ecology approach facilitates sustainable plant protection by identifying odorant receptors (ORs) tuned to odorants, especially the volatile molecules emitted from host plants that insects use for detection. A few studies have explored such an approach to develop sustainable pest management programs, especially in host-specialized insect species. We revealed the molecular mechanism of host plant detection of a destructive, invasive insect pest of palm trees (Arecaceae), the Asian palm weevil (*Rhynchophorus ferrugineus*), by deorphanizing an OR (RferOR2) tuned to several palm-emitted odors. We found that RferOR2 responded explicitly to several ecologically relevant palm-emitted odors and significantly to palm esters when transgenically expressed in *Drosophila* olfactory neurons. We mapped RferOR2 expression in the *R. ferrugineus* genome and found that odor specificity is likely to develop equally in both sexes. We inferred that the semiochemicals that attract palm weevils to a palm tree might aid in weevil control efforts by improving attraction, enticing many palm weevils to the traps. We demonstrate that including synthetic palm volatiles in pheromone-based mass trapping has a synergistic effect on pheromones, resulting in significantly increased weevil catches. We proved that insect OR deorphanization could aid in the identification of novel behaviorally active volatiles for inclusion in pest management. These results suggest that targeting RferOR2 may help design receptor antagonists that can interfere with weevil host-searching behavior in sustainable pest management applications.

**Significance:** Asian and South American palm weevils are tremendously important agricultural pests primarily adapted to palm trees and cause severe destruction, threatening sustainable palm cultivation worldwide. The host plant selection of these weevils is mainly attributed to functional specialization of odorant receptors that detect palm-derived volatiles. We unraveled the intricacies of weevil–palm tree communication by deorphanizing an odorant receptor tuned to natural palm-emitted odors. We used palm ester volatiles, which produced a significant response in the functional studies, and proved their synergistic effect on the pheromone coinciding with increased weevil catches in the field. We revealed that insect odorant receptor deorphanization could help identify novel behaviorally active volatiles (reverse chemical ecology) for sustainable palm protection.

## Introduction

Palm trees of the family Arecaceae are cultivated worldwide for commercial (e.g., coconut, date, oil, and ornamental palms), world heritage (e.g., The Palmeral of Elche, Spain), and cultural icon (e.g., Cannes in southern France) purposes. The United Nations Food and Agriculture Organization (FAO) acknowledged palm trees, such as date palms, as agricultural crops closely connected with human life, and hence, they were included on UNESCO’s list of the Intangible Cultural Heritage of Humanity (https://ich.unesco.org/en/RL/date-palm-knowledge-skills-traditions-and-practices-01902). The Asian palm weevil, *Rhynchophorus ferrugineus* (also known as the red palm weevil, RPW), which is endemic to Southeast Asian countries, has been documented since the mid-last centuries, and it is now considered the world’s most destructive quarantine category-1 pest, spread throughout palm-growing areas worldwide (Supplementary, Fig. S1). This invasive pest causes slow palm tree death and almost always remains unnoticed until the tree falls ^1^ (Fig. 1). During the last three decades, this destructive weevil moved beyond its native range (Southeast Asian countries), where its preferred host was coconut (*Cocos nucifera* L.), and invaded date palms (*Phoenix dactylifera* L.) in Middle Eastern countries (see Fig. 1) and Canary Island date palm (*Phoenix canariensis* Wildpret) in Mediterranean countries, accelerating its risk to palm agriculture ^2^ (Supplementary, Fig. S1). Similarly, the RPW’s American counterpart, the South American palm weevil (SAPW), *R. palmarum*, which is native to South America, has posed a destructive threat to palm trees, affecting coconut and oil palm (*Elaeis guineensis* Jacq.) production in America (Supplementary, Fig. S1). In addition, *R. palmarum* is a vector of a plant pathogenic nematode, the red ring nematode, that causes the lethal palm disorder red ring disease, amplifying the threat to ornamental and date palms in California ^3^. Both palm weevil invasions have caused significant damage to palm trees, triggering environmental, economic, and cultural threats to palm agriculture. Several insecticides are used indiscriminately to combat the severe threat caused by this devastating insect pest, with severe adverse impacts on the environment and human health, in addition to the development of resistance to insecticides in the weevils ^4^. To address these concerns, many palm-growing countries seek alternatives and more sustainable palm protection against this weevil.

**Fig. 1.**
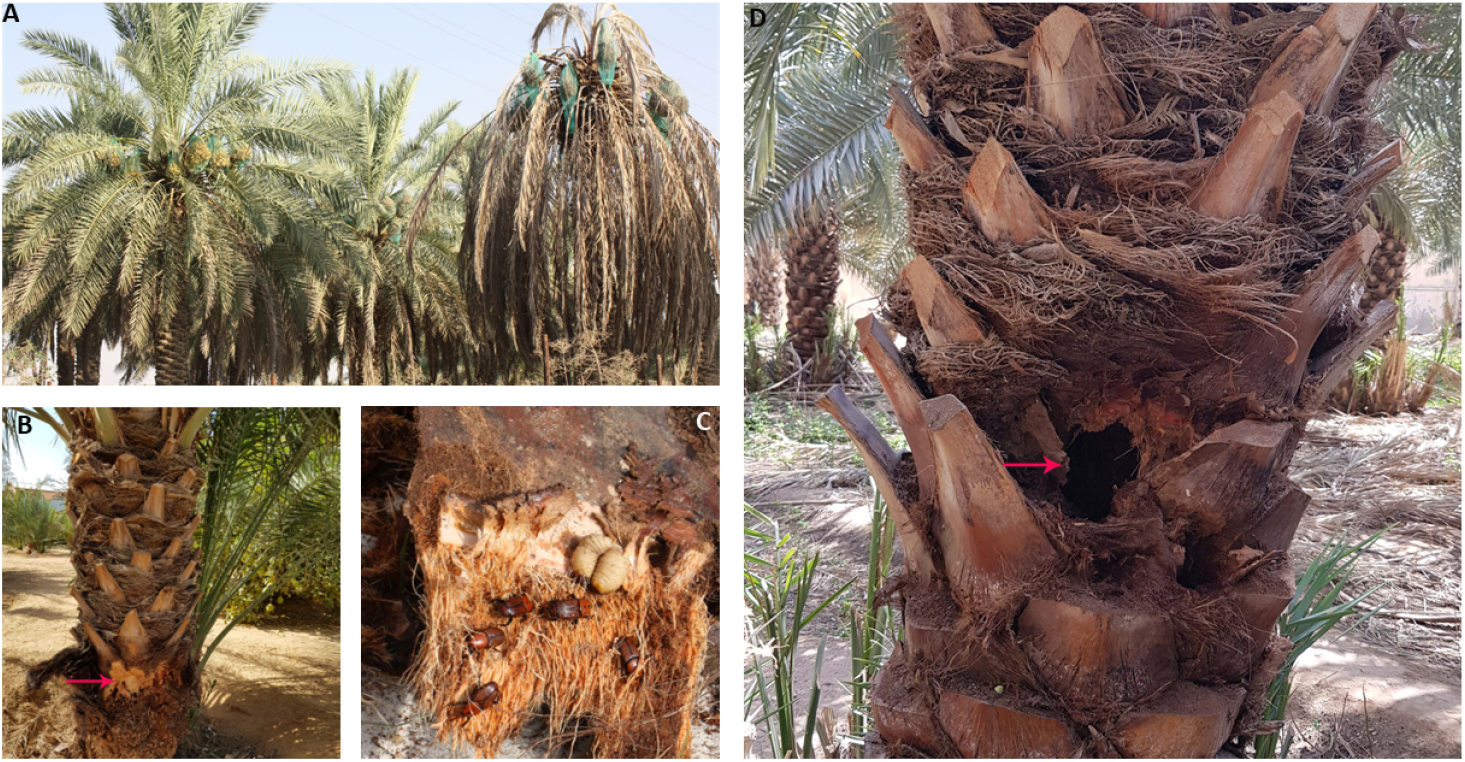
Red palm weevil (*Rhynchophorus ferrugineus*) infested date palm (*Phoenix dactylifera* L.) trees (Al Kharj, Saudi Arabia). **A**. drying of infested offshoots (right); **B**. oozing of brownish fluid together with palm tissue (arrow), **C**. presence of adults and grubs at the base of fronds; and **D**. typical tunneling of a palm tree by grubs (arrow).

Palm weevils rely on chemical communication for mate finding and host searching ^5-7^. Several behavioral studies and field trap catches using specific palm-derived volatiles have indicated that palm esters elicit the most robust behavioral responses when used in traps as coattractants and pheromone synergists ^8-10^. The active compounds identified through conventional chemical ecology approaches are being used worldwide based on bioassay-electrophysiology-guided results, which often leads to unsatisfactory weevil control ^2^. Recent advancements in understanding the molecular basis of olfaction, especially uncovering host–insect communication through functional studies of odorant receptors (ORs), have helped identify key host volatiles that mediate host selection and paved the way for the emergence of reverse chemical ecology ^11-13^. Insect chemosensory systems detect odors (olfaction) through olfactory sensilla residing in the antennae where olfactory receptor neurons (ORNs) express ORs ^14^. A receptor unit is composed of a unique ‘tuning’ OR that recognizes a set of odorants and a coreceptor (Orco) that forms heteromeric complexes with tuning receptors to enable sensory signaling ^15,16^. As the OR confers chemical sensitivity to the heteromeric receptor complex, it has an essential role in discriminating an enormous variety of odorants ^17^. The host-emitted volatile organic compounds (VOCs) identified through reverse chemical ecology approaches may be used in pest management programs, particularly in pheromone-based trapping to enhance pest catches. Such reverse chemical ecology may facilitate sustainable plant protection by including volatile molecules as attractants or synergists in mass-trapping of insect pests ^12^. We applied reverse chemical ecology approaches to unravel the complexities of weevil–palm tree communication and identified the palm-derived compounds eliciting the most activity in the palm weevil. These active palm-derived compounds were identified by their interaction with palm weevil OR when transgenically expressed in *Drosophila* and proved to be efficient synergists or coattractants in the mass trapping of palm weevils in the field. This study contributes to knowledge of chemical communication in the *R. ferrugineus* through functional characterization of palm ester-specific OR and their genome-level organization. It exemplifies the efficiency of reverse chemical ecology for sustainable pest management applications.

## Methods

### Red palm weevil rearing and *Drosophila* stocks

The RPWs used in this study originated from a laboratory culture and the field. RPWs were maintained on sugarcane stems as described previously ^7^. The laboratory-reared RPW culture was considered pure-line, as there was no mixing with other populations. Field RPW populations (males and females separately) were collected from Al Qassim (25.8275° N, 42.8638° E) in Saudi Arabia (see details in Supplementary Methods).

Transgenic *Drosophila melanogaster* Or67d^GAL4 18^ and *UAS-RferOR2* lines were maintained on a yeast diet, as previously reported ^19^.

### Chemicals, pheromones, and insecticide

Twenty-nine chemical compounds, including several palm-derived volatiles (Supplementary, Table S1), synthetic palm esters, commercial pheromone, Ferrolure^+TM^, thirty-two pheromone compounds (Supplementary, Table S2), and an insecticide, Carbofuran (Sumo® 3% CG), were used in this study (see details in Supplementary Methods).

### RPW transcriptome, OR expression mapping, and differential gene expression analysis

Illumina HiSeq 2000 sequencing was performed at the core sequencing facility of KAUST, Saudi Arabia (see details in Supplementary Methods). Briefly, the RPW antennal and snout (rostrum) transcriptomes were generated using laboratory-reared (pure-line) and field-collected adult male and female samples. Total RNA was extracted from ten pairs of antennae from either male or female *R. ferrugineus* (laboratory-reared and field-collected) using a PureLink RNA Mini Kit (Invitrogen, USA). The quantity and quality of the total RNA were validated using a Qubit 2.0 Fluorometer (Invitrogen, Life Technologies), and RNA integrity was further confirmed using a 2100 Bioanalyzer (Agilent Technologies). Following the manufacturer’s protocols, paired-end cDNA libraries were prepared using the TruSeq Stranded mRNA preparation Kit (Illumina Inc.). The cleaned RNA-seq reads were mapped to the *R. ferrugineus* genome ^20^ (GenBank assembly accession: GCA_014462685.1) using the ‘RNA-seq analysis procedure’ implemented in CLC Workbench /Server suite (Qiagen). Gene expression levels were quantified and reported as reads per kilobase of transcript per million mapped reads (RPKM) and transcripts per kilobase of exon model per million mapped reads (TPM).

We visualized the expression level of the RferORs in comparison with RferOrco ^21^, with two other functionally characterized ORs, *viz*., RferOR1 ^7^ and RferOR41 (previously RefOR6) ^22^ in the CLC Genomics Server. Differential expression analysis was conducted in the CLC Genomics Server. The differential expression level of all genes was calculated by the tool ‘Differential expression in two groups’ for male vs. female and field vs. laboratory groups (see details in Supplementary Methods). The expression levels of the transcripts were expressed as normalized TPM values of RferOR mRNA using male vs. female and laboratory vs. field *R. ferrugineus* transcriptomes (see details in Supplementary Methods).

### RNA extraction, cloning of the full-length RferOR2 gene, and sequence analysis

We used previously annotated *R. ferrugineus* OR sequences ^7,23,24^ for oligonucleotide design (Supplementary, Table S3). Total RNA was extracted from 30 mg of antennae tissue of laboratory-reared male and female RPW adults using the PureLink RNA Mini Kit (Thermo Fisher, Waltham, MA, USA). To obtain the full-length open reading frame (ORF) of ORs, we amplified the 5′ and 3′ cDNA ends using the rapid amplification of cDNA ends (RACE) technique (see details in Supplementary Methods).

### Functional study of RferOR2

#### Transgenic expression of RferOR2 in *Drosophila* ORNs

We followed previously described methods for cloning the *RferOR2* ORF into the *pUAST*.*attB* vector ^7^. Briefly, the following *RferOR2* gene-specific primers were designed: 5′-GAATTCATGAAGCCTGTCAAGTATCGTGAATTG-3′ (an *EcoR*I site is included) and 5′-GCGGCCGCCTATATAGATAACTGATTTTCTGATGC-3′ (a *Not*I site is included). First-strand cDNA was synthesized using SuperScript IV Reverse Transcriptase (Thermo Fisher) from 1 µg of total RNA extracted (PureLink RNA Mini Kit, Thermo Fisher) from 10-day-old male antennae (see details in Supplementary Methods). Transgenic *D. melanogaster UAS*-*RferOR* lines were generated by BestGene Inc. (Chino Hills, CA, USA) by injecting the EndoFree *pUAST*.*attB*-*RferOR2* plasmid into fly embryos expressing the PhiC31 integrase and carrying an *attP* landing site within the ZH-51C region of the second chromosome ^25^. *Drosophila* lines expressing the RferOR2 transgene in at1 ORNs (genotype *w*; *UAS-RferOR2, w*^+^; *Or67d*^GAL4^) were generated by crossing the *UAS*-*RferOR2* line with the *Or67d*^GAL4^ line ^18^. Genomic integration and expression of *RferOR2* in *Drosophila* were verified using PCR and RT‒PCR on *Drosophila* genomic DNA and antennal RNA, respectively (Supplementary, Table S3).

#### Single-sensillum recordings and odor simulation

We used 2- to 5-day-old *RferOR2*-expressing *Drosophila* flies for single-sensillum recordings (SSRs) on at1 sensilla by following standard procedures ^7,26^. The flies were kept alive under a constant 1.5 L.min^-1^ flush of charcoal-filtered air; humidified air was delivered to the antenna until the recording finished. A wide range of palm volatiles and behavior-modifying compounds (Table S1) and coleopteran pheromones (Supplementary, Table S2), including *R. ferrugineus, R. palmarum*, and *R. phoenicis* pheromones, dissolved in mineral oil were first tested at 100 µg on filter paper to draw response spectra of *Drosophila* ORNs expressing *RferOR2*. After that, we performed a dose‒ response analysis of highly active palm esters with increasing doses of 0.001, 0.01, 0.1, 1.0, 10, and 100 µg on clean filter paper strips (see details in Supplementary Methods). Responses to the different stimuli were compared to the response to solvent alone using one-way analysis of variance (ANOVA) followed by a post hoc Tukey honestly significant difference (HSD) test (Bonferroni α-level at 0.05) using statistical calculators (https://astatsa.com/).

### Phylogenetic analysis, gene structure, and conserved motif analysis

Phylogenetic analysis was performed using OR amino acid sequences from *R. ferrugineus* ^23^, *R. palmarum* ^24^, *Ips typographus* ^27^, *Megacyllene caryae* ^28^, and *Nicrophorus vespilloides* ^29^. Multiple sequence alignments were performed using MAFFT v.7 ^30^, with the auto (FFT-NS-1, FFT-NS-2, FFT-NS-i, or L-INS-i; depending on data size) strategy and default parameters, followed by manual trimming to remove gaps and ambiguous sequences. We used the auto algorithm and BLOSUM62 as the scoring matrix. The final multiple sequence alignment contained 350 sequences with 1126 amino acid sites used in the IQ-tree phylogenetic tree ^31^. The automatic model search was performed using ModelFinder ^32^, and the JTT+F+I+G4 substitution model was determined as the best-fitting model according to the Bayesian information criterion (BIC). The maximum likelihood analysis was performed using default settings and ultrafast bootstrap support ^33^ with 1000 replicates in IQ-tree ^31^ (see details in Supplementary Methods).

The RferOR sequences were correctly annotated and mapped to the *R. ferrugineus* genome ^20^ (GenBank accession numbers GCA_014462685.1 and GCA_014490705.1) using a BLASTN search against the *R. ferrugineus* genome created on Geneious *v*7.1.9 (Biomatters) and correctly annotated. To identify the antenna-specific RferORs ^7^, we generated expression profiles in the male and female antennae (from the laboratory and field). We compared them to male and female snout (rostrum) (from the laboratory and field) transcriptomes. The *R. ferrugineus* snout raw reads were deposited in the NCBI Sequence Read Archive. The assembled and cleaned *RferOR2* reads in the antennae and snout transcriptomes were visualized with the CLC Genomics Server, using marked gene positions in the *R. ferrugineus* genome (GenBank: GCA_014462685) ^20^ (see details in Supplementary Methods).

### Structural modeling and docking of RferOR2

Modeling of the 3-dimensional (3D) structure of RferOR2, identification of the potential binding pockets in the protein, and docking screening with the target analytes were carried out as reported in ^7,34^. We then conducted RferOR2 docking predictions with a common insect repellent, N,N-diethyl-3-methylbenzamide (DEET) (CAS 134-62-3) (53), and different palm ester volatiles to determine *in silico* interactions and modes of action with RferOR2. The docking predictions were viewed and analyzed using the Swissdock server plugin UCSF Chimera (52).

### Reverse chemical ecology for palm weevil management

#### RPW field trap catches with synthetic palm esters

We selected three palm esters found to elicit high activity of RferOR2 *Drosophila*, viz., ethyl propionate (EP), ethyl isobutyrate (EI), and ethyl acetate (EA), which were used as coattractants for the RPW pheromone trapping field experiments. The experiment was conducted on a coconut plantation at the farm (15.10600 N, 74.14860 E) of Sulcorna village, Quepem town, Goa, India, from June to October 2022 (temperature, 24-30 °C, RH 80-90%, rainfall 2600-2800 mm). The coconut plantation in the experimental plot was approximately 2 ha, containing approximately 200 coconut trees (*Cocos nucifera* L., variety Benaulim), and was reported to be moderately infested with *R. ferrugineus* by the local agriculture department. Standardized pheromone traps and field trials were designed and carried out as previously described ^2^. For all field trials, we used a commercial pheromone lure (Ferrolure^+TM^, ChemTica International, San Jose, Costa Rica, purity > 98% with a 4-10 mg/d release rate). Each treatment comprised pheromone (Ferrolure^+ TM^), fresh sugarcane food bait, and a synthetic palm ester or blend. The food bait was mixed in the water with 1 g of carbofuran to kill captured weevils and prevent escape (see details in Supplementary Methods). A total of twelve treatment combinations with seven replications of each trial were performed.

We normalized the RPW catch data using log10 (*y*+1) transformation ^35^ and analyzed them using one-way ANOVA, and means were compared using post hoc Tukey’s HSD tests (see details in Supplementary Methods).

## Results

### RPW transcriptome assemblies, OR expression mapping, and differential gene expression analysis

Antennal and snout transcriptome data (field and laboratory RPWs) were uploaded to the NCBI under BioProject PRJNA275430 and SRA accession numbers, *viz*., antennae: SRR22098129; SRR22098128; SRR22098127 and SRR22098126; and snout: SRR17732029, SRR17732028, SRR17732027 and SRR17732026. *De novo* transcriptomes were assembled for both male and female antennae and snout (field and laboratory) (see Supplementary, Table S4). BUSCO analysis was performed separately on each male and female transcriptome. It resulted in hits for 98.27% of queried sequences for both transcriptomes, and 91.9% and 89.6% were identified as complete in the male and female transcriptomes, respectively.

We annotated antennal RferORs based on TPM values and mapped their expression level distribution obtained in the transcript quantification of the male and female transcriptome and laboratory-reared RPWs versus field-collected RPWs compared to that of the pheromone receptor (RferOR1) ^7^ and OR coreceptor, RferOrco ^21^ (Fig. 2A). The results indicated that OR expression in both males and females was more similar than the samples from different conditions (laboratory vs. field) (Fig. 2A). RferOrco was highly expressed (both in males and females) under laboratory and field conditions, followed by RferOR1 and RferOR2 (Fig. 2A). RferOR2 was the second most highly expressed antenna-specific OR among 69 ORs expressed in the antennae (Fig. 2A). Furthermore, RPW genome-wide analysis and expression profiling of male and female adult antennae (both laboratory and field conditions) confirmed the highest expression of RferOR1 and RferOR2 (Supplementary, Tables S5 and S6). In addition, based on the normalized expression, RferOR1 and RferOR2 were the first and second most highly expressed *R. ferrugineus* ORs, respectively, in both male and female adult antennae and were found to be upregulated under laboratory conditions (Fig. 2A) (Supplementary, Table S5, and S6). A previous functional study revealed RferOR1 as a pheromone receptor narrowly tuned to the RPW aggregation pheromone ^7^. Thus, we focused on RferOR2, as this is the second most highly expressed antenna-specific OR, and we also focused on RferORs belonging to the same phylogenetic clade (see Supplementary, Fig. S3). The expression values of the other three ORs in the RferOR2 clade (RferOR7, RferOR41, and RferOR44) were meager compared to those of RferOR1 and RferOR2 in both male and female adult antennae, indicating the important role of the latter two ORs in *R. ferrugineus* chemical communication (Supplementary, Tables S5 and S6). We calculated the percentage of expression increase or decrease (relative expression) in RferOR2-clade genes relative to RferOR1. The results showed a 15.31% and 19.54% (male and female, respectively) decrease in RferOR2 expression compared to RferOR1 expression. In contrast, other ORs in the RferOR2 clade showed more than a 70% decrease in their expression in RPW antennae. Nevertheless, based on the expression values and fold changes in expression, RferOR2 expression patterns between male and female antennae were not significantly different; however, they were slightly higher in males and thus did not show sex-biased expression (Supplementary, Tables S7 and S8) (FDR *p value*: 0.999990485 and Bonferroni: 1). We generated the RferOR2 expression profile in the antennae and snout transcriptome by mapping the gene position in the *R. ferrugineus* genome (GenBank: GCA_014462685). The results revealed strict antenna-specific expression of RferOR2, as we did not find the same in the snout transcriptome (Fig. 2B). In general, we observed that RferOR2 was the best OR that needs to be functionally characterized, as this OR is antenna-specific (Fig. 2B) and highly expressed (Fig. 2A; Supplementary, Tables S5 and S6); hence, we proceeded with cloning and transgenic expression in *Drosophila* ORNs.

**Fig. 2.**
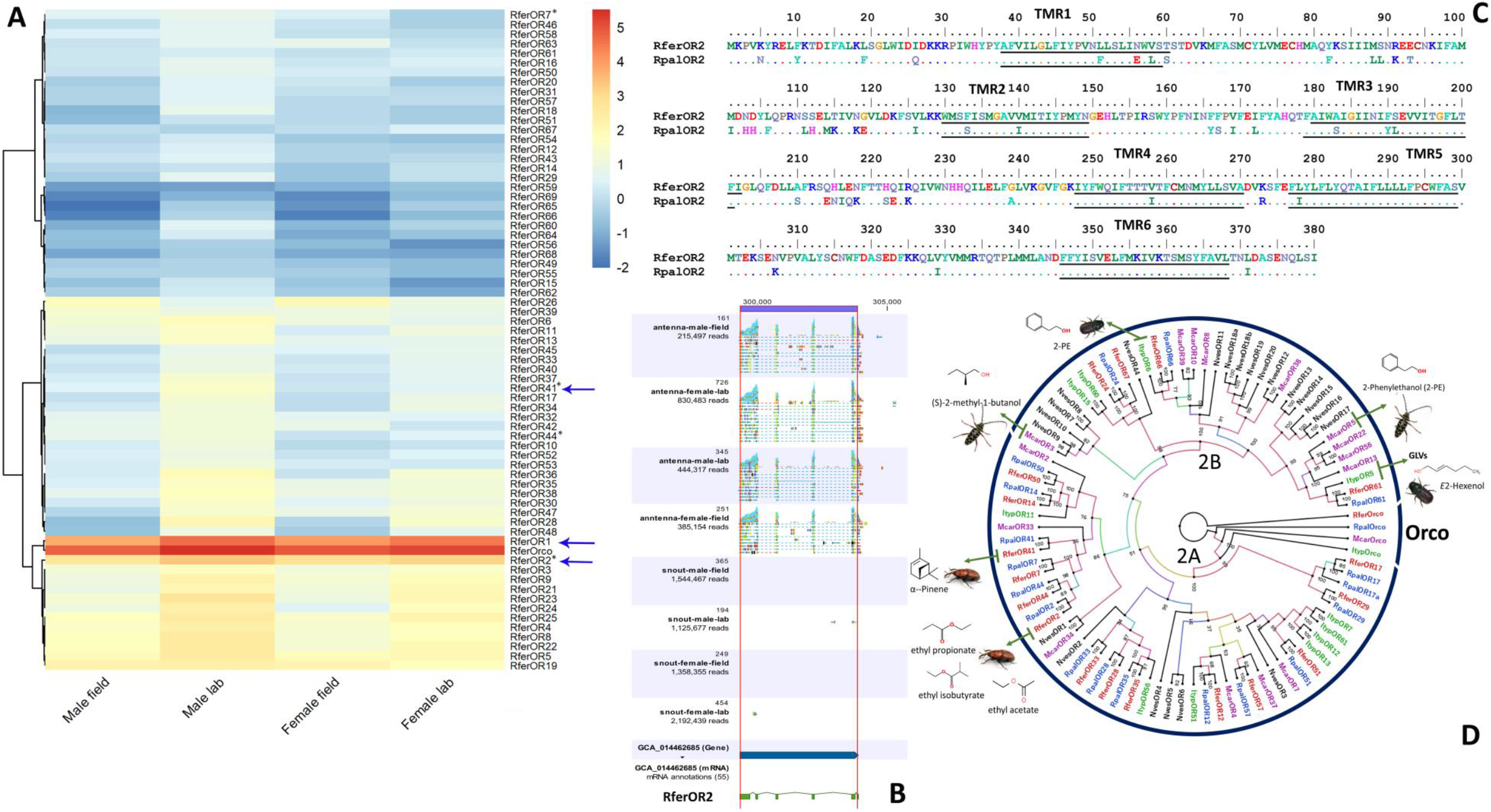
**A** Normalized expression profile of *R. ferrugineus* odorant receptors (RferORs) in the antennae of male and female (laboratory-reared and field-collected). Expression levels of RferOR1 to RferOR69 and RferOrco of the aforesaid antennal transcriptomes, represented as heat plot based on the log2-transformed transcripts per million (TPM) values. The lightest blue color represents the lowest expression. * Indicate the ORs from the RferOR2 clade. The deorphanized RferORs are shown with blue arrows. **1B**. RferOR2 expression profile in the male and female antennae and snout (laboratory and field) transcriptomes with marked gene position on the RPW genome (GenBank: GCA_014462685. **1C**. Deduced amino acid sequence of *R. ferrugineus* OR2 (RferOR2) aligned with the ortholog RpalOR2 of *R. palmarum*. Identical residues are denoted by dots. The transmembrane regions (TMRs) are underlined. **ID**. Phylogeny of the group 2A and 2B subfamilies of coleopteran odorant receptors (ORs), rooted with the conserved Orco lineage. The tree was built from the MAFFT alignment of OR amino acid sequences of *R. ferrugineus*, Rfer (red), *R. palmarum*, Rpal (blue), and the following coleopteran species: *Nicrophorus vespilloides*, Nves (black); *Ips typographus*, Ityp (green); and *Megacyllene caryae*, Mcar (purple). Functionally characterized ORs (see Supplementary, Table S12) in group 2A and 2B subfamilies and their ligand molecules are shown. Numbers on the branches are bootstrap values (UFBoot *n* = 1000). Scale = 2.0 amino acid substitutions per site. The branch appearance was colored based on the bootstrap values.

### RferOR2 cloning and sequence analysis

We obtained an ORF of the RferOR2 cDNA transcript containing 1143 bp, corresponding to a protein of 380 amino acids (aa) with a predicted molecular weight of 44.72 kilodaltons (Supplementary, Table S9). We analyzed the RferOR2 gene in the GenBank database and retrieved the RferOR2 ortholog from a closely related species, the American counterpart, *R. palmarum* (RpalOR2) ^24^. RferOR2 shares 83.42% identity with RpalOR2. The multiple sequence alignment of RferOR2 and RpalOR2 shows highly conserved C-terminal parts (amino acids 226-380) (Fig. 2C). Seven canonical transmembrane domains (pfam02949) were identified in RferOR2 and RpalOR2 and were predicted to be located in the plasma membrane, which is a typical characteristic of insect ORs (Fig. 2C; Supplementary, Table S9). The conserved protein domain family of RferOR2, seven-transmembrane (7tm_6) G-protein-coupled receptor class (sensory perception of smell, GO:0007608), was predicted to be positioned at 69-362 (63-362 for RpalOR2) based on SMART domain architecture analysis (e-value 1.8e-38) and an NCBI conserved protein domain (accession, cl20237) search ^36^. Based on blastx searches, RferOR2 shares 49.03% identity with *Sitophilus oryzae* odorant receptor-4 (GenBank acc no. XP_030746544.1). Sequence comparison of RferOR2 with other coleopteran counterparts revealed that it belongs to the group 2 subfamily of the coleopteran ORs (Fig. 2D). Some ORs in the group 2 subfamily are responsible for host plant volatile detection (see Fig. 2D; Supplementary, Fig. S3).

### Functional study of RferOR2

#### RferOR2 responds to palm esters and structurally related compounds

We successfully expressed RferOR2 specifically in the at1 sensilla of *Drosophila* (*w; UAS-RferOR2; Or67d*^Gal4^). We used SSRs to record the responses of the flies to 100 µg of twenty-nine ecologically relevant palm ester, ketone and alcohol compounds (Supplementary, Table S1). The SSR results showed that RferOR2-expressing *Drosophila* ORNs exhibited the strongest response to several palm esters, viz., ethyl propionate, ethyl isobutyrate, ethyl butyrate, ethyl acetate, methyl butyrate, methyl isobutyrate, propyl butyrate, 2-pentanone, acetoin, 2-butyl acetate, and isobutyl propionate (Fig. 3A). Among the palm esters tested, ethyl propionate elicited the strongest response (mean response of 152 spikes/s) in the screening experiments (7500% increased action potential compared to that for solvent alone) (Tukey HSD Q = 10.62), followed by slightly weaker and similar responses to ethyl acetate (122 spikes/s) (Q = 8.51), ethyl isobutyrate (114 spikes/s; Q = 7.95), and ethyl butyrate (109 spikes/s; Q = 7.59) (F = 16.41; df = 4,49; *P* < 0.00001) and an even weaker response to 2-phenyl ethanol (19 spikes/s; Q = 3.81), 1-butanol (15 spikes/s; Q = 3.00), 3-buten-2-ol (12 spikes/s; Q = 2.24) and ethyl valerate (7 spikes/s; Q = 1.25) (F = 2.22; df = 4,49; *P* < 0.082). In addition, RferOR2 *Drosophila* responded to 2-pentanone, acetoin, 2-butyl acetate, and isobutyl propionate with overall weaker or lower response magnitudes (Fig. 3A). Interestingly, RferOR2 *Drosophila* ORNs showed a slight response to 2-phenyl ethanol (Fig. 3A), an ecologically relevant compound of the conifer-feeding curculionids *I. typographus* and *D. ponderosae* ^37^.

**Fig. 3.**
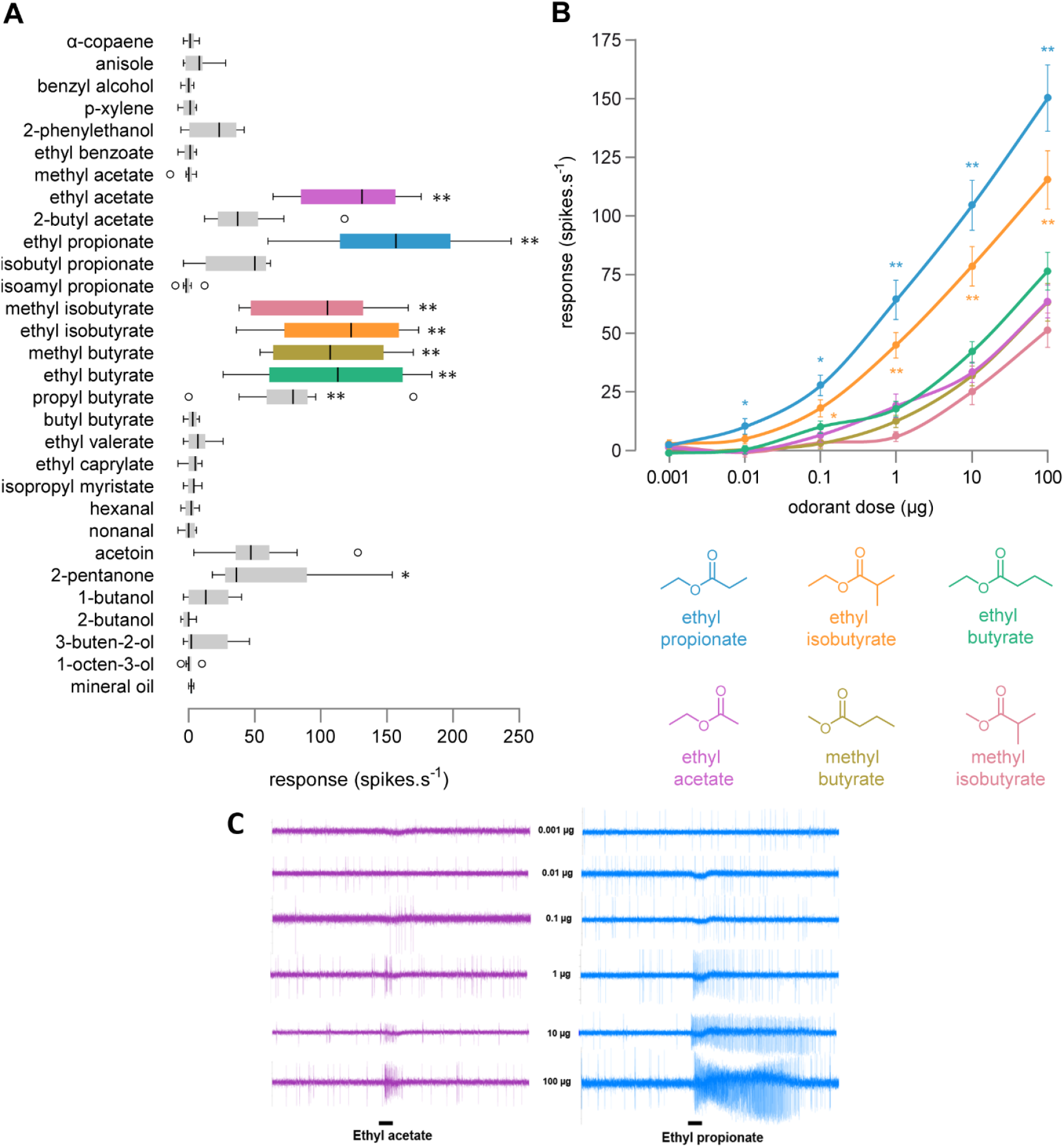
**A** RferOR2, when expressed in *Drosophila* ORNs, is activated by several palm esters, ketone and alcohol compounds. Action potential frequency of *Drosophila* at1 ORNs expressing *RferOR2* when stimulated with a panel of palm ester compounds and related chemicals (100 µg loaded in the stimulus cartridge). Box plots show the number of spikes/s ±SEM (*n* = 10). Significance levels are denoted at ** *P*<0.01 and * *P*<0.05 in comparisons to solvent (mineral oil) alone (ANOVA followed by Tukey HSD test, Bonferroni and Holm multiple comparison tests). **3B**. Dose-dependent responses of RferOR2 expressed in *Drosophila* at1 ORNs to the six most active palm ester compounds. Data represented are mean action potential frequencies ± SEM (*n* = 14-15). At 100 µg, ethyl propionate and ethyl isobutyrate show the highest responses, followed by other palm ester compounds. Significance levels are denoted at ** *P*<0.01 and * *P*<0.05 for multiple comparisons (ANOVA followed by Tukey HSD test, Bonferroni and Holm tests). **3C**. Representative single sensillum recordings were obtained for a RferOR2 *Drosophila* at1 ORNs stimulated with increasing doses of ethyl acetate and ethyl propionate. Black bars represent the duration of the stimulus (100 ms).

We next recorded the dose-dependent response curve (dose of 0.001 µg to 100 µg of stimulus within the stimulus cartridge) of RferOR2 *Drosophila* ORNs with the six most active palm esters. At the lowest dose (0.001 µg), the *p* value corresponding to the F-statistic of one-way ANOVA was higher than 0.05, suggesting that the 0.001 µg dose did not induce a response, regardless of the compound (F = 1.96; df = 5, 89; *P* = 0.0929). At the 0.01 µg dose, ethyl propionate exhibited a significantly greater response than ethyl butyrate and methyl butyrate (*P* < 0.05) and ethyl acetate and methyl isobutyrate (*P* < 0.01). RferOR2 *Drosophila* ORNs exhibited significant dose-dependent responses to ethyl propionate and ethyl isobutyrate (Fig. 3B), indicating that RferOR2 preferentially binds to these two esters. At higher doses (0.1 to 100 µg), ethyl propionate and ethyl isobutyrate elicited significantly stronger responses than other compounds, with a *p* value lower than 0.05 (*P* < 0.01 and *P* < 0.001) (Fig. 3B). For each dose, responses to ethyl propionate and ethyl isobutyrate were not significantly different (F = 0.27; df = 1,11; *P* < 0.61) (Fig. 3B), indicating that RferOR2 binds these molecules equally well when heterologously expressed in *Drosophila* ORNs. RferOR2 showed slightly lower but similar sensitivities to other active palm esters, *viz*., ethyl butyrate, ethyl acetate, methyl butyrate, and ethyl isobutyrate ((Fig. 3C). These four compounds activated RferOR2 in a dose-dependent manner (Fig. 3B), although the response curves were significantly weaker than those of ethyl propionate and ethyl isobutyrate.

#### RferOR2 failed to respond to RPW aggregation pheromone and structurally related compounds

To confirm whether RferOR2 is exclusively tuned to palm esters and palm-derived compounds, we performed SSR recordings with RferOR2 *Drosophila* to a wide range of palm weevil and beetle aggregation pheromone compounds (Supplementary, Table S2). The SSR results showed that RferOR2 *Drosophila* flies failed to respond to any pheromone compounds, including ferrugineol, ferrugineone, rhynchophorol, phoenicol, and structurally related compounds (Supplementary, Fig. S4).

### Phylogenetic analysis, gene structure, and conserved motif analysis

OR sequences obtained from publicly available transcriptome and genome projects of various Coleoptera species were used to build the phylogenetic tree. RferORs were clustered in four (1, 2, 5, and 7) of the seven major OR groups, as previously reported ^7^ (Supplementary, Fig. S3), and RferOR2 and RpalOR2 were clustered in major coleopteran OR group 2 (Fig. 2D). Group 2 was split into two subfamilies (groups 2A and 2B) ^29^, and the RferOR2 clade was distributed in the group 2A subfamilies (Fig. 2D). Interestingly, this clade also contains a nonpalm plant volatile (α-pinene)-tuned OR, RferOR41 (GenBank acc no. MW979236) (Fig. 2D), identified recently in *R. ferrugineus* ^22^, which shared 32.86% amino acid identity with RferOR2 (Table S10). Within this clade, two orthologous RferORs (RferOR44 and RferOR7) were positioned close to RferOR2 (Fig. 2D), with amino acid identities of 38.56% and 39.21%, respectively (Table S10). The *R. ferrugineus* (RferOR1) ^7^ and *I. typographus* ^27^ pheromone receptors were located in coleopteran OR group 7 (Fig. S3). Based on the function and distribution of ORs among the 2A and 2B subfamilies, one could predict that ORs from group 2 are involved in host plant volatile detection ^27^ (Fig. 2D and Supplementary, Fig. S3).

Using NCBI DBSOURCE accession JAACXV010014020.1 locus_tag=“GWI33_016023” (299313..303900) (scaffold_65774), we annotated a deduced amino acid sequence of the RferOR2 fragment that was identical to *R. ferrugineus* RferOR2. The functional RferOR2 gene length mapped was 4588 bp in scaffold_65774 (Supplementary, Fig. S5). The genomic organization of RferORs within the RferOR2 clade revealed that they are distributed across different scaffolds in the *R. ferrugineus* genome with an uneven distribution pattern (Supplementary, Fig. S5). The pheromone receptor (RferOR1) ^7^ and odorant coreceptor (RferOrco) ^21^ were mapped to scaffolds 63 and 235, respectively (Supplementary, Fig. S5). To characterize the structural diversity of the RferORs in the RferOR2 clade, their intron‒exon organization was analyzed. Basic information about these RferOR reads mapped in the male and female transcriptomes with the ratio of unique exon and intron reads is provided in Supplementary, Table S5, and Table S6. The largest RferOR2 exon and intron comprised 659 bp and 1418 bp, respectively. The majority of ORs in the RferOR2 clade contained five exons, except RferOR41, which possessed six exons when compared to the *R. ferrugineus* pheromone receptor (RferOR1) and odorant coreceptor (RferOrco), which possessed eight and eleven exons, respectively (Supplementary, Tables S5 and S6). The largest intron (>6 kb) was found in the clade with RferOR7, followed by RferOR41 (>5 kb) and RferOR44 (<4 kb) (Supplementary, Fig. S5). The exon length, intron number, and intron phase were highly conserved within the same gene group of male and female *R. ferrugineus* (Supplementary, Tables S5 and S6).

The MEME motif analysis conducted on the RferOR2 clade (RferOR2, OR7, OR41, and OR44) revealed a similar motif consensus distribution pattern (Supplementary, Fig. S6) that diverged from that of RferOR1 and RferOrco (Supplementary, Fig. S6). All the RferOR2 clade proteins, including RpalOR2, contained multiple transmembrane regions and were relatively conserved (Supplementary, Fig. S2; Table S9). These ORs were predicted to show an intracellular N-terminus membrane orientation consistent with the known insect OR membrane topology.

### Structural modeling and molecular docking of RferOR2

The 1143-bp ORF of RferOR2 encodes a protein of 380 amino acid residues that were modeled (Fig. 4). We identified a total of seventy-four pockets (with a minimum size of 1.4 Å), with a main binding pocket (the predicted active site) (Fig. 4A) made up of 34 amino acid residues (Supplementary, Table S11), of which twenty-four were hydrophobic (70.59%), five were hydrophilic (14.71%), four were positive (11.46%), and two were negative (5.88%). Compared to the data obtained in SSR, in which ethyl propionate and ethyl isobutyrate induced significantly greater responses than ethyl acetate, RferOR2 docking showed a slightly higher affinity to ethyl propionate and ethyl isobutyrate (−21.22 kcal/mol and −23.11 kcal/mol, respectively) than to ethyl acetate (binding energy, −17.03 kcal/mol). Within the active site, 2 (out of the 34, Table S11) residues (Y160 and F173) showed direct interactions with ethyl propionate (69 total interactions), ethyl acetate (5/34: Y72, I142, Y160, F173, and Y279) (117 total interactions) or ethyl isobutyrate (3/34: I142, Y160, and F173) (116 total interactions). The common residues for all three ligands were I142, Y160 and F173. Residues Y72 and Y279 interacted with ethyl acetate, while none interacted with ethyl propionate and ethyl isobutyrate. We then conducted docking studies using the well-known insect-repellent DEET (Fig. 4D and E). The binding energy value was used to define the RferOR2 binding affinity to DEET; the lower the binding energy is, the stronger the binding (higher affinity), and *vice versa*. The docking results showed that DEET exhibited higher affinity to RferOR2, with an interaction energy of −30.67 kcal/mol. The results revealed that DEET bound to RferOR2 with higher affinity than palm esters. DEET bound to the RferOR2 main binding pocket by interacting with five residues, T154, R157, S158, N163, and I164 (Fig. 4E).

**Fig. 4.**
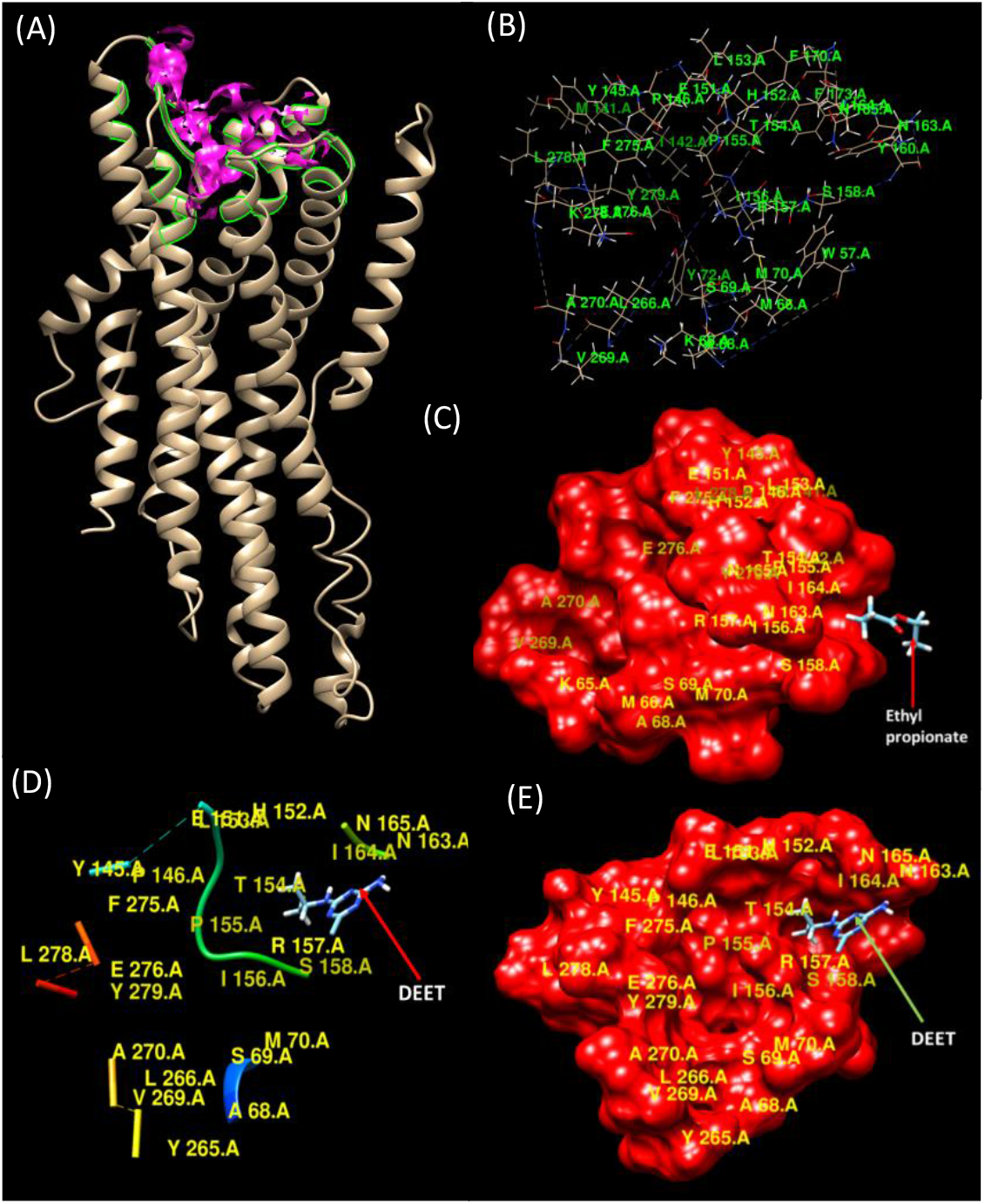
**(A)** Cartoon representation of Helix and Loop structure of RferOR2. The active site is highlighted in pink within the whole protein, and the constituent amino acid residues are labelled. In **(B)** and **(C)**, the pocket is shown on its own, and the constituent amino acid residues are labelled in both figures. In **(C)**, the pocket is shown in hydrophobicity surface representation and colored in red. DEET binds to the binding pocket of RferOR2 **(D** and **E)**. In **(E)**, the pocket is shown in hydrophobicity surface representation and colored in dark red.

### Reverse chemical ecology for palm weevil management

#### RPW field trap catches with synthetic palm esters

A total of 536 adult weevils were captured during June-October 2022 experimental trials. The total adult catches observed using widely evaluated pheromone traps [lure, food bait, plus ethyl acetate (EA) ^2^] were marginally higher (statistically nonsignificant; F = 0.0201; df = 2, 20; *P* < 0.98) (Tukey HSD results, *P* < 0.89) than those observed when using the other two coattractant [ethyl propionate (EP) and ethyl isobutyrate (EI)] traps for the mean (±) of 4-week field trials (Fig. 5). In addition, in the Bonferroni and Holm simultaneous comparison, relative to treatment EA, the other treatments had nonsignificant effects (Bonferroni and Holm *P value*, EA vs. EP = 1.72, and EA vs. EI = 1.99 and 0.99, respectively). Enhanced adult weevil catches were recorded when we used a pheromone trap with a blend of 3:2:1 (EA:EP:EI or EP:EA:EI), followed by a blend of 100:1 (EA:EP) and 30:1 (EA:EP), which were significantly different (*P*<0.01 and *P*<0.05 Tukey HSD results) from the pheromone trap with EA (Fig. 5). Similarly, a larger number of adult RPW catches was obtained in the pheromone traps with coattractant blends of 30:1 and 50:1 (EA:EP) compared to EA, EP or EI alone in the pheromone traps (Fig. 5). Compared to conventional pheromone traps with lure plus EA alone, the most significant adult RPW catches were recorded for a coattractant blend of 3:2:1 [(EA:EP:EI) or (EP:EA:EI)], supported by Bonferroni and Holm values (significant at *P*<0.01), followed by 100:1 (EA:EP) (Bonferroni and Holm *P*<0.05 and *P*<0.01). Nevertheless, when compared to the traps complemented with lure plus EA alone, the traps complemented with EA and EP (30:1, 50:1, and 100:1) and EP:EA:EI at 3:2:1 showed enhanced weevil catches (F = 3.98; df = 4, 34; *P* < 0.05). We calculated the percentage of increase or decrease in adult weevil catches in each treatment compared to the pheromone trap with EA alone (Fig. 5). The results showed 246.02%, 187.74%, 173.54%, 153.48%, and 124.51% increases in the RPW adult catches with a synthetic palm ester blend of 3:2:1 (EA:EP:EI), 3:2:1 (EP:EA:EI), 100:1, 30:1 and 50:1 (EA:EP), respectively, over the course of 28 days (Fig. 5).

**Fig. 5.**
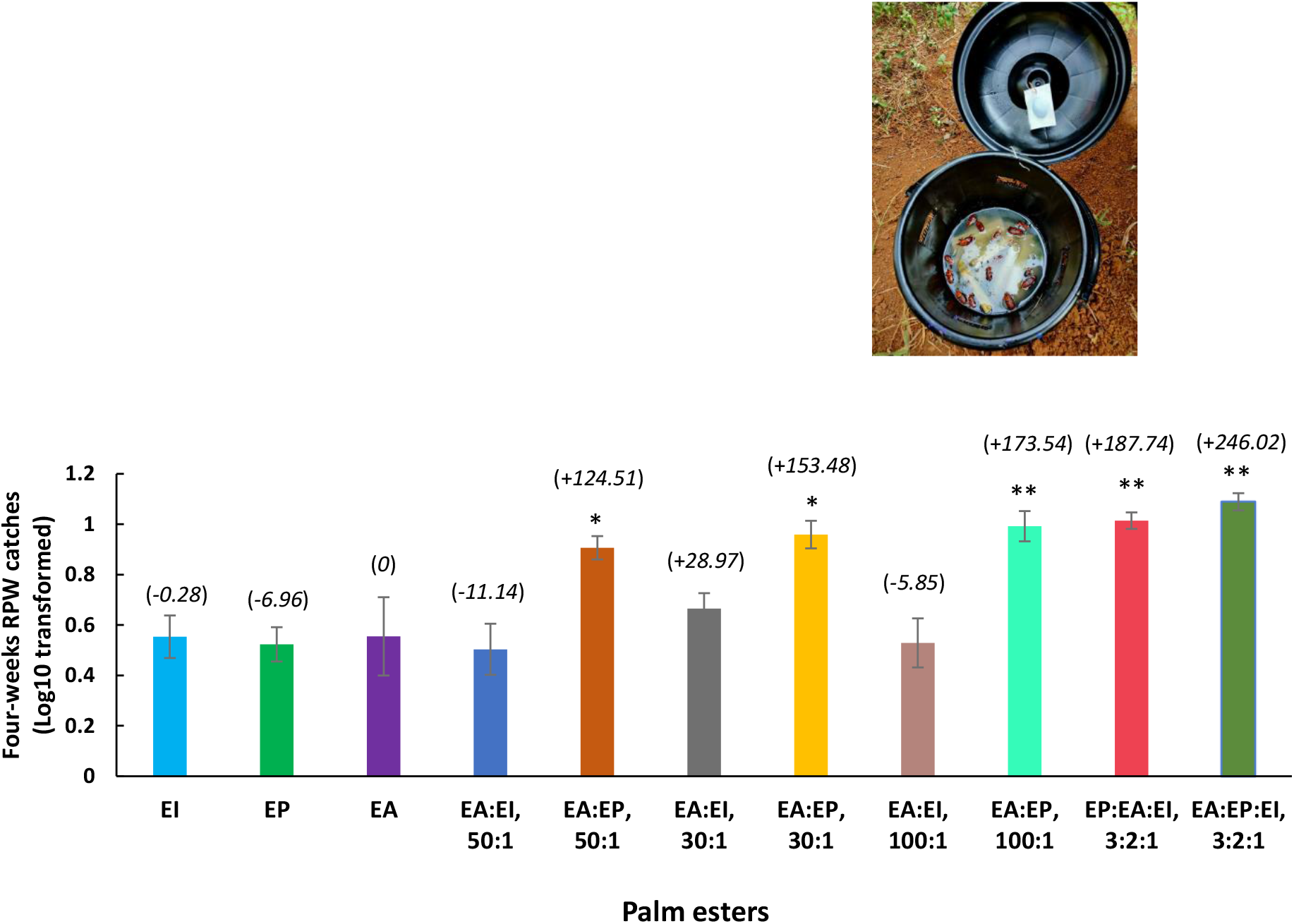
Field effectiveness of palm esters in mass trapping of *R. ferrugineus* in a coconut farm. Mean (± standard error of the mean) numbers of four weeks (June to October 2022) RPW adult catches were normalized using log10 (*y*+1) transformation (see details in *Supplementary Methods*). *N*=7 for each treatment group. Traps were baited with the aggregation pheromone (Ferrolure^+TM^), sugarcane food bait and supplemented with the following synthetic palm esters; ethyl propionate (EP), ethyl isobutyrate (EI), and ethyl acetate (EA), both single compound and in the blend. Asterisks indicate significance levels at * *P*<0.05 and ** *P*<0.01 (ANOVA followed by Tukey HSD test and Bonferroni and Holm multiple comparison tests, compared to each pair relative to EA). Numbers in parenthesis over each bar are back-transformed (*Supplementary Methods*) percentages of adult catches that increased (+ sign) or decreased (−sign) in the traps relative to EA. The inset photo shows *R. ferrugineus* adult caches in the pheromone trap.

## Discussion

There is growing concern over the invasion, range expansion, and rapid spread of the RPW *Rhynchophorus ferrugineus* and the South American palm weevil *R. palmarum* worldwide, which have resulted in significant and accelerating risks to sustainable palm agriculture (Supplementary, Fig. S1). The rate of human-mediated translocation of insect pests such as palm weevils has increased noticeably in the last century ^38,39^. Information on the functional divergence of the OR gene family in palm weevils may be essential in understanding their host adaptation, range expansion, and global spread. Furthermore, understanding how palm weevils detect their host’s volatile emissions—particularly as they spread across new areas and colonize host plants such as date palms and ornamental palms—could help us design sustainable pest control strategies. With this vision, we functionally characterized an *R. ferrugineus* OR (RferOR2) that is broadly tuned to natural palm-emitted odors, so-called “palm esters,” when expressed in transgenic *Drosophila* fruit flies. The occurrence of an OR broadly tuned to natural palm-emitted odors suggests that this OR is involved in the palm weevil’s adaptive radiation on various palm trees. We show that reverse chemical ecology may facilitate sustainable plant protection by including volatile molecules identified in the weevil–palm tree interactions for the mass-trapping of palm weevils.

Initially, we hypothesized that palm-emitted volatiles would be detected by a highly expressed OR from the group 2 subfamily of coleopteran ORs (Supplementary, Fig. S3) ^7^, as this group includes the most previously reported plant volatile ORs ^37,40-42^. To test this hypothesis, we selected the antenna-specific (Fig. 2B) and second most highly expressed OR from group 2–RferOR2–and successfully expressed it in *Drosophila* at1 neurons. We tested RferOR2 with an array of palm-derived VOCs and confirmed that transgenic fly at1 ORNs responded to these compounds noticeably, with high sensitivity to ethyl propionate, ethyl isobutyrate, ethyl butyrate, methyl butyrate, and ethyl acetate. Interestingly, RferOR2 *Drosophila* could not detect any pheromone compounds tested, including *R. ferrugineus, R. phoenicis*, and *R. palmarum* aggregation pheromones. This result was consistent with those of several previous functional studies of coleopteran ORs in the group 2 subfamily that revealed that these ORs were tuned to host plant volatiles ^37,40-42^. In contrast, ORs in the group 7 subfamily were tuned to pheromone compounds (Supplementary, Fig. S3) ^7,27^.

To date, few studies have focused on the volatile detection of host plants in Coleoptera. Specifically, only thirteen ORs from six different coleopteran insects have been functionally characterized (see Supplementary, Table S12). Palm weevils belong to Curculionidae, the largest family of beetles, with more than 80,000 species that primarily rely on chemical communication ^43^. Palm weevils, *Rhynchophorus* spp., are attracted explicitly to wounded, stressed, or fermented palms, particularly around tunnel openings and other damaged areas ^2,44,45^. Ethyl esters are common volatile constituents in fermenting palm oils, sap, and plant tissues, and many of these esters elicit significant electrophysiological responses in palm weevil species ^8-10,46-48^. Our studies revealed that RferOR2 responded to several palm esters, ketones (2-pentanone and acetoin), and alcohols (2-phenyl ethanol), which are all reported as key components of date palm ^49-51^, coconut ^52,53^, and Canary Island palm ^47^. Often, RPW traps are baited with fermenting palm material and/or date fruits that act synergistically with the aggregation pheromone lure of the weevils ^54^ as a result of RPW’s ability to detect palm esters, as demonstrated in the current study. In addition, ethyl acetate is a common volatile reported in almost all palm trees found to influence palm weevil behaviors ^9,55,56^. We showed that complementing pheromone–food bait traps with ethyl acetate leads to a slightly higher number of RPW catches than using ethyl propionate (Fig. 5). Nevertheless, the use of ethyl propionate, ethyl acetate, ethyl isobutyrate and their combinations with the pheromone lure and food bait substantially and significantly increased the RPW catches, which may result from the broadly tuned RferOR2 response to palm esters (Fig. 5). As RferOR2 detected all these compounds, the activity of this OR might play an important role in this synergistic attraction. Hence, it would be highly recommended to use palm ester combinations as a pheromone synergist or coattractant for mass trapping of palm weevils. Similarly, such a strategy may also be successful with the South American palm weevil, *R. palmarum*, and the African palm weevil, *R. phoenicis*, as several studies have proven antennal detection and behavioral response to palm esters ^8,46,56^. In addition, studies have demonstrated that the release of the *R. palmarum* aggregation pheromone starts approximately 10 minutes after the insect detects ethyl acetate and continues for several hours, highlighting the importance of RferOR2 in palm weevil ecology and behavior ^57^. These findings support the hypothesis that RferOR2 in palm weevils evolves with a genus-specific olfactory function that predominantly detects palm esters, which have both behavioral and ecological relevance, as this receptor conveys the message of a nonhost plant that should be avoided. *R. ferrugineus* and *R. palmarum* behavioral and electrophysiology studies demonstrated antennal detection of palm esters ^8,9,46,47,55,56^. The functional characterization of OR2 orthologues in diverse *Rhynchophorus* species is needed to confirm whether OR2 has undergone functional conservation in conveying the message of the host plant.

Looking at the OR phylogenetic tree, we observed that RferOR2 and its ortholog RpalOR2 are members of a monophyletic clade that clusters with three other *R. ferrugineus* ORs, RferOR41, RferOR7, and RferOR44 (Fig. 2D). Intriguingly, RferOR2 shares high bootstrap support with RferOR44, which has been previously reported to be ubiquitously expressed ^7^. Our previous study indicated that RferOR7 has antenna-specific expression, and RferOR41 was also expressed in legs, in addition to antennae ^7^. The RferOR2 clade genes were mapped to different scaffolds, including RferOR44, a close relative of RferOR2, for which a genome-wide analysis identified the complete gene (Supplementary, Fig. S5). The gene structure conservation, the moderately high amino acid sequence homology, and common motif sites in the deduced proteins suggest that at least RferOR2, RferOR7, and RferOR44 originated from gene duplication (Supplementary, Fig. S3), a frequent mechanism observed in insect OR evolution and acquisition of new detection capacities ^58,59^. The RferOR2 protein shared a single motif with the pheromone receptor RferOR1 (WVYHWTD) (Supplementary, Fig. S6), indicating a high degree of sequence divergence and supporting their functional separation, as we demonstrated that RferOR1 ^7^ and RferOR2 (current study) detect RPW pheromone and palm volatiles, respectively. In addition, a recent study revealed the functional role of RferOR41 in detecting the nonhost plant volatile α-pinene, which causes a significant avoidance response in the RPW ^22^. Functional studies also revealed that RferOR41 could not detect any palm esters, including ethyl acetate ^22^, indicating that the members of the RferOR2 clade have divergent functions. These findings support the hypothesis that within the same OR clade, OR2 in palm weevils evolves with a specific olfactory function that predominantly detects palm esters, whereas OR41 conveys the message of a nonhost plant that should be avoided. Revealing the response spectrum of the remaining ORs (RferOR44 and RferOR7) will help understanding the OR functional evolutionary history in detecting palm esters or other volatile compounds.

Apart palm esters, we investigated whether RferOR2 detects other palm volatiles by including two key ketones (2-pentanone and acetoin) commonly reported as palm volatiles ^49-51^. Our results showed that RferOR2 did not exclusively detect palm esters (Fig. 3A), as a weak response to 2-pentanone and acetoin (Fig. 3A) could be observed. Acetoin is a major naturally occurring date palm volatile ^51^; hence, detection of this compound is vital for palm weevil host localization and colonization. The capacity of RferOR2 to detect palm esters and other palm host volatiles, such as acetoin, may have facilitated rapid diversification and range expansion of the palm weevil. *R. ferrugineus* is reportedly native to southeast Asia and Melanesia, where its original host is coconut ^38^. Nevertheless, in the last century, the weevil expanded its range through multiple anthropogenic introductions and became a pest of date and Canary Island palms ^2,38^. Unraveling the electrophysiological response of RferOR2 to key VOCs of coconut, oil, Canary Island palms, and date palms will help understand its role in the weevil expansion.

Interestingly, the RferOR2 *Drosophila* ORNs responded with slightly weak sensitivity to 2-phenyl ethanol (2-PE), an ecologically relevant volatile reported for several conifer-feeding bark beetles, pine weevils, and cerambycid beetles ^28,37^ (Table S12). 2-PE has been reported to be part of an attractive odor bouquet released by the fungal symbionts of the bark beetle *Ips typographus* ^60^ and has also been reported to be produced by the mountain pine beetle *Dendroctonus ponderosea*, in which 2-PE acts as an aggregation pheromone antagonist ^37^. Additionally, 2-PE was found to be produced by the gut bacteria of the pine weevil, *Hylobius abietis*, and it acts as a strong antifeedant and deterrent ^37^. The ORs tuned to 2-PE have recently been identified from all three species of weevils from the Curculionidae family (ItypOR6/DponOR8/HabiOR3), and all belong to the coleopteran OR subfamily 2B ^37^. We found an ortholog in *R. ferrugineus* and *R. palmarum* (RferOR66/RpalOR66) with high bootstrap support (Fig. S3), and it could be hypothesized to have a similar function in detecting 2-PE. Interestingly, our OR phylogeny indicated that RferOR2 belongs to the coleopteran OR subfamily 2A (Fig. 2D; Supplementary, Fig. S3), which is distantly related to ItypOR6/DponOR8/HabiOR3. 2-PE has already been reported in palms ^51^. It would be more interesting to know whether 2-PE is an antifeedant/deterrent in other curculionids and, thus, why RferOR2 detects it, which is tuned to attractants. Further functional studies are needed to confirm that RferOR66/RpalOR66 detects 2-PE, as RferOR2 does, and to hypothesize on possible adaptive radiation of OR functions in Curculionidae.

Finally, we inferred from the RferOR2 docking studies that among palm esters, ethyl propionate, ethyl isobutyrate, and ethyl acetate showed higher affinities, representing the best ligands for RferOR2. Interestingly, the common insect-repellant DEET exhibited a significantly higher affinity toward RferOR2 than any palm esters used in this study (Fig. 4). By being the best ligand for RferOR2, together with the information available regarding the DEET inhibitory function on other insect ORNs ^61,62^, we can hypothesize that the interaction of DEET with RferOR2 may produce an inhibitory effect on palm-emitted odor detection. It likely disrupts weevil–palm tree communication. Nevertheless, more functional studies and behavioral bioassays must be performed to confirm the above hypothesis. Our docking data provide essential information on RferOR2 residues that interact with DEET, which might be useful for additional investigation. If successful, DEET may interfere with the detection of palm esters and may challenge the extended effects of palm weevil host-seeking behavior.

## Conclusion

RferOR1 and RferOR2 are essential for pheromone communication (within species) and host plant volatile detection (host searching) and play an important role in *R. ferrugineus* fitness and survival. Maintaining a high degree of specificity in pheromone detection through the narrowly tuned receptor RferOR1 ^7^ maintains high fidelity in the mate recognition system, ensuring reproductive success ^63^. Equally noteworthy, a broadly tuned receptor (RferOR2, current study) would give palm weevils an advantage in detecting diverse volatiles from their food source (palms), which would help them find ovipositional sites and colonies. The functional study of RferOR2 led us to propose new pheromone synergistic candidates, which proved effective in field experiments and increased palm weevil catches. Such a so-called reverse chemical ecology approach paves the way for discovering new synergistic compounds. Adopting the newly proposed blends or including ethyl propionate with ethyl acetate as pheromone synergists in the mass trapping of palm weevils worldwide has a great potential in pest management applications. As RferOR1 and RferOR2 detect ecologically relevant compounds, targeting these ORs for the development of receptor blockers or antagonists or for genome editing could disrupt the chemical communication of the quarantine pest. Their discovery, thus, enables substantial novel approaches for a sustainable way of controlling palm weevils.

## Acknowledgments

The authors extend their appreciation to the Deanship of Scientific Research, King Saud University, for funding through the Vice Deanship of Scientific Research Chairs, the chair of date palm research. This work was financially supported through research grants from King Abdullah University of Science and Technology (KAUST) in Saudi Arabia (KAUST-OSR-2018-RPW-3816-1, OSR-2018-RPW-3816-4 and BAS/1/1020-01-01). We thank Cam Oehlschlager (ChemTica International, San Jose, Costa Rica) for generously providing the Ferrolure^+TM^ and palm ester emulsions and for his expert advice on field trials. The authors also thank the date palm farmers in the Al-Qassim areas for their support in obtaining red palm weevils.

## Authors’ contributions

BA, EJ.-J, KP, and AP conceived of the study and acquired the grant. BA, RS, and MAA participated in RPW field collection and rearing. JJ, BA, SM, and AP performed transcriptomics, assembly, and annotation. NM, AC, and EJ.-J performed the *Drosophila* transgenic expression and electrophysiology work. NM, AC, BA, and EJ.-J analyzed the SSR results. RS and BA carried out field experiments and analyzed the data. KC, KP, and BA performed docking studies. JJ and BA performed RPW genome reassembly, odorant receptor annotation, and analysis. BA drafted the manuscript, and all authors contributed to, edited, and approved the final version.

## Competing interests

BA, NM, EJJ, and AP are inventors of palm ester combinations in the field trap experiment and may have a financial interest in patent applications related to RPW pest management. BA, NM, EJJ, KP, KC, and AP have interests in filing patent applications related to *R. ferrugineus* RferOR2-biosensor for pest management applications. The remaining authors declare that they have no competing financial interests.

## Supplementary information

### Supplementary Methods

**Fig. S1** A graphical illustration of *Rhynchophorus ferrugineus* and *R. palmarum* distributions worldwide.

**Fig. S2** Predicted transmembrane regions of RferOR2/RpalOR2 clade proteins by HMMTOP, TMHMM and Phobius.

**Fig. S3** Maximum likelihood consensus tree of ORs from Coleoptera.

**Fig. S4** RferOR2-expressing *Drosophila* failed to respond to the RPW aggregation pheromone and structurally related compounds.

**Fig. S5** Comparison of the scaffold distribution of *R. ferrugineus* RferOR2 clade genes.

**Fig. S6** MEME motif analysis performed on the RferOR2 clade proteins.

**Table S1** List of molecules tested on RferOR2 *Drosophila* via SSR.

**Table S2**. List of aggregation pheromone compounds (with the corresponding species) and structurally related chemicals used to stimulate ORNs expressing RferOR2.

**Table S3**. List of primers used for RferOR expression studies.

**Table S4**. Red palm weevil antennal and snout transcriptome assembly report.

**Table S5**. Genome-wide analysis and expression profiling of antenna-specific ORs (RferOR1-RferOR22) and ORs from the RferOR2 clade in the RPW male antennal transcriptome.

**Table S6**. Genome-wide analysis and expression profiling of antenna-specific ORs (RferOR1-RferOR22) and ORs from the RferOR2 clade in the RPW female antennal transcriptome.

**Table S7**. Expression level distribution of RferORs from the RferOR2 clade, RferOR1, and RferOrco, obtained from the transcript quantification of male and female RPWs from the field vs. laboratory.

**Table S8**. Mapping the expression level distribution of RferORs from the RferOR2 clade, RferOR1, and RferOrco, obtained from the transcript quantification of RPWs from the field and laboratory - male vs. female.

**Table S9** Identification and characteristics of *R. ferrugineus* RferOR2 clade genes in comparison to the pheromone receptor (RferOR1).

**Table S10**. Pairwise amino acid identity generated based on MAFFT multiple alignments.

**Table S11**. List of the amino acid residues of the RferOR2 active site.

**Table S12**. List of coleopteran ORs functionally characterized and detecting host/plant volatiles.

## Data availability

The RPW antennal and snout transcriptomes (male and female; laboratory and field) have been submitted to the NCBI SRA SUB12148378 and SUB10967960 under BioProject PRJNA275430.

## Notes

### Competing Interest Statement

The authors have declared no competing interest.

