## Supplementary for "Reverse chemical ecology approach for sustainable palm tree protection against invasive palm weevils"

[1]*King Saud University, Chair of Date Palm Research, Center for Chemical Ecology and Functional Genomics, Department of Plant Protection, College of Food and Agricultural Sciences, Riyadh 11451, Saudi Arabia.*
[2]*INRAE, Sorbonne Université, CNRS, IRD, UPEC, Université Paris Cité, Institute of Ecology and Environmental Sciences of Paris, iEES-Paris, 78000 Versailles, France.*
[3]*King Abdullah University of Science and Technology (KAUST), Bioscience Programme, BESE Division, Thuwal, Jeddah 23955-6900, Saudi Arabia.*
[4]*University of Ljubljana, Biotechnical Faculty, Agronomy Department, SI-1000 Ljubljana, Slovenia.*
[5]*Don Bosco College of Agriculture, Agricultural Entomology Department, Sulcorna, Goa 403705, India.*
[6]*The University of Manchester, Department of Chemical Engineering, Manchester, M13 9PL, UK.*

**This PDF file includes:**

Supplementary Methods

Figures S1 to S6

Tables S1 to S12

SI References

### Supplementary Methods

#### Red palm weevil rearing and *Drosophila* stocks

Red palm weevils used in this study originated from a laboratory culture and the fields. The laboratory culture was established in 2009. Since then, weevils have been maintained on sugarcane stems at 28–30°C with a photoperiod of 18 h:6 h (light: dark cycle), ~ 60% RH as described previously (1). Lab-reared RPW culture was considered pure-line, as there was no mix with other populations. Field RPW populations (male and female separately) were collected from Al Qassim (25.8275° N, 42.8638° E) in Saudi Arabia from date palm orchards where high infestations have been reported. The adult male and female antennal dissections were performed after insects were anaesthetized using $CO_2$ for 1-2 min.

#### Chemicals, pheromone and insecticide

Twenty-nine chemical compounds, including several palm ester volatiles (Table S1), synthetic palm esters, commercial pheromone, Ferrolure$^{+TM}$, and thirty-two pheromone compounds (Table S2), including RPW and SAPW aggregation pheromones, (4*RS*,5*RS*)-4-methylnonan-5-ol (ferrugineol) and 4(*RS*)-methylnonan-5-one (ferrugineone), and *4S-(E)-6-Methyl-2-hepten-4-ol* (rhynchophorol), and African palm weevil (APW) aggregation pheromone, 3-Methyloctan-4-ol (phoenicol) were purchased [(>98% purity), ChemTica International, San Jose, Costa Rica].

Carbofuran (Sumo® 3 % CG), which was purchased from Kakode farm, Margao (Goa, India), is a suspension concentrate containing 4% w/w calcium silicate, 5% w/w blank granules, and a 91% w/w g.a.i./L carbofuran (a 3% concentration of the active ingredient, carbofuran).

#### RPW transcriptome, OR expression mapping, and differential gene expression analysis

Image deconvolution and quality value calculations were performed using Illumina GAPipeline1.3. Illumina adaptors were detected and removed by an automatic read-through adapter trimming option implemented in the 'Trim Reads' tool of Qiagen CLC Genomics Server (CLC) (*v* 21.0.1). Low-quality bases were also trimmed off, allowing two ambiguities per read. Filtered paired-end reads were QC validated through a 'QC for Sequencing Reads Tool' of CLC. A reference de novo transcriptome assembly was constructed with 'De Novo Assembly Tool' of CLC and contigs functionally annotated by the BLAST2GO command line tool (*v*1.5).

The criteria of significant differential expression were |log2 fold change |≥ 1 (2), False Discovery Rate (FDR) ≤ 0.001(3), and Bonferroni post hoc analysis. Differentially expressed ORs (up-and downregulated) in male and female antennae under lab and field conditions were plotted in heatmaps which were created using pheatmap (*v*1.0.12) R package (4). The resulting *P*-values were corrected using Benjamini and Hochberg's method for managing the FDR. The OR genes with a corrected *P*-value < 0.05 found by pheatmap were considered as differentially expressed genes.

To compute the distances between expression values of genes, we used the squared Euclidean distance measure, and the clustering of genes was performed using Ward.D2 agglomeration method. A value of 1 was added to the TPM value of each gene before the log2 transformation to avoid infinite values. Bonferroni post hoc analysis was estimated across different tissues (male vs. female and lab vs. field), and hierarchical clustering was performed using Multi Experiment Viewer (MeV *v* 4.9.0).

#### RNA extraction, cloning of full-length RferOR2 gene, and sequence analysis

The cDNAs were prepared from *R. ferrugineus* adult male, and female antennae total RNA (~ 1 μg) using a SMARTer RACE Kit (Clontech, Mountain View, CA, USA). Touchdown polymerase chain reaction with gene-specific primers (Table S3) and universal primer mix [95 °C for 5 min, 35 cycles of 95 °C for 1 min, 65 °C (touchdown to 55 °C) for 30 s, and 72 °C for 3 min; and one cycle at 72 °C for 10 min] was carried out using an Advantage 2 PCR kit (Clontech), and the PCR products were gel-purified (Wizard SV Gel purification kit; Promega, Madison, WI, USA) cloned into the pGEM-T vector (Promega) followed by transformation into JM109 competent cells (Promega). The plasmids were isolated manually and sequenced in both directions (ABI 3500, Thermo Fisher) for sequence verification. Sequences were aligned and annotated using a BLASTx homology search.

The transmembrane helices were predicted using TMHMM-2.0 (https://services.healthtech.dtu.dk/service.php?TMHMM-2.0), Phobius (https://phobius.sbc.su.se/) and HMMTOP (http://www.enzim.hu/hmmtop/). Signal peptides and subcellular localization prediction were performed by SignalP5.0 (https://services.healthtech.dtu.dk/service.php?SignalP-5.0 /). Molecular weight (MW) and isoelectric point (pI) were predicted using ExPASy (http://web.expasy.org/protparam/). Conserved domains were analyzed by SMART (http://smart.embl-heidelberg.de/) and CD-search, with the default parameters (http://www.ncbi.nlm.nih.gov/Structure/cdd/ docs/cdd_search. html). The pairwise identity matrix was generated by SIAS (http://imed.med.ucm.es/Tools/sias.html).

#### Functional study of RferOR2

#### Transgenic expression of RferOR2 in *Drosophila* ORNs.

PCR was performed by using Applied Biosystems ProFlex PCR System (Thermo Fisher), the conditions of which had 94°C for 5 min, 35 cycles at 94°C for 15 sec, 60°C for 30 sec, and 72°C for 3 min, and a final extension at 72°C for 10 min. The PCR product was separated by agarose gel electrophoresis and purified from the agarose gel with a Wizard® SV Gel and PCR Clean-up system (Promega). The purified DNA was ligated into a pGEM®-T easy vector and transformed into JM109 *E. coli* cells (Promega). Plasmid DNA (pGEMT-RferOR2) was purified using a QIAprep Miniprep kit (Qiagen, Venlo, Netherlands), and the insert was sequenced (ABI 3700, Thermos). The linearized plasmid DNA was double digested with *EcoR*I and *Not*I (New England Biolabs, Ipswich, MA, USA), separated by agarose gel electrophoresis, and purified with the Wizard® SV Gel and PCR Clean-up system. The purified RferOR2 insert was ligated into the linearized *pUAST.attB* vector, which had been predigested with *EcoR*I and *Not*I, and the recombinant *pUAST.attB-RferOR2* plasmid was transformed into TOP10 *E. coli* cells (Thermo Fisher). The *pUAST.attB-RferOR2* plasmid construct was purified using the EndoFree Plasmid Maxi kit (Qiagen), then sequenced.

**Single-sensillum recordings and odor simulation.**

The response of at1 neurons to each stimulus was recorded using an EX-1 amplifier (Dagan, Minneapolis, MN, USA), a Digidata 1440A acquisition board (Molecular Devices, Sunnyvale, CA, USA), and analyzed using the pCLAMP 10 software (Molecular Devices). Cartridges containing 10 µg of 11-*cis*-vaccenyl acetate (cVA, the ligand of the *Drosophila* receptor OR67d) or mineral oil only were used as controls. Responses of at1 ORNs expressing RferOR2 were calculated by subtracting the spontaneous firing rate (measured 500 ms before the stimulation) from the firing rate during the stimulation.

#### Phylogenetic analysis, gene structure, and conserved motif analysis

The tree was rooted with the Orco lineage, visualized and edited with FigTree *v*1.4 (tree.bio.ed.ac.uk), colored, and finally edited with Adobe Illustrator (Adobe, CA, USA).

We mapped the exon-intron positions in the genome at the scaffold region (299313-303900) flanking in the locus_tag="GWI33_016023 (NCBI acc. JAACXV010014020.1) (5) were aligned with RferOR2 5' UTR, coding sequence, 3' UTR using MAFFT program *v*7 (6). To illustrate the gene structure, the exon-intron structure, including exon positions and gene length, was constructed manually. Multiple Expectation for Motif Elicitation (MEME) (http://meme-suite.org/tools/meme) were used to predict the protein motifs (parameters: number of repetitions: any, the maximum number of motifs: 10, optimum motif width set to > 6 and < 200). The predicted MEME motifs were searched in the Expasy-Prosite database with the ScanProsite server (https://prosite.expasy.org/scanprosite/).

#### RPW field trap catches with synthetic palm esters

These synthetic palm esters (emulsions of 100 mg/day optimized release rate) dispensers used in this study were made explicitly for the field trials at the research facility at ChemTica International (San Jose, Costa Rica).

Pheromone traps were set on the ground under the shade of a palm canopy about 20 m apart in each block. Traps were allowed to remain at a given spot in the field for seven days and moved sequentially from one site to another every week, ensuring that each treatment was placed at every location in the area for seven days (to neutralize treatment bias, if any, due to spot effect). Pheromone traps were serviced once a week when the bucket traps', food bait, water, and insecticide were renewed.

We conducted post-hoc Tukey HSD Test with all field trails were simultaneously compared, and each trail relative to pheromone lure, food bait plus ethyl acetate caches using Bonferroni and Holm multiple comparison tests (https://astatsa.com/). The increase or decrease in the total count of RPW adult catches was calculated using back-transformed data represented as a percentage of increase or decrease relative to pheromone lure plus ethyl acetate (for example, [(lure, EP – lure, EA) / |lure, EP|] × 100). The back-transformed data in percentages were generated (52), and significant mean differences were accepted at the 0.05% probability level.

**Fig. S1** A graphical illustration of *Rhynchophorus ferrugineus* and *R. palmarum* distributions worldwide (based on the CABI invasive species compendium, https://doi.org/10.5061/dryad.m93f6) (last accessed on September 27, 2022) on commercially important palms (coconut, date, oil, and ornamental palms).

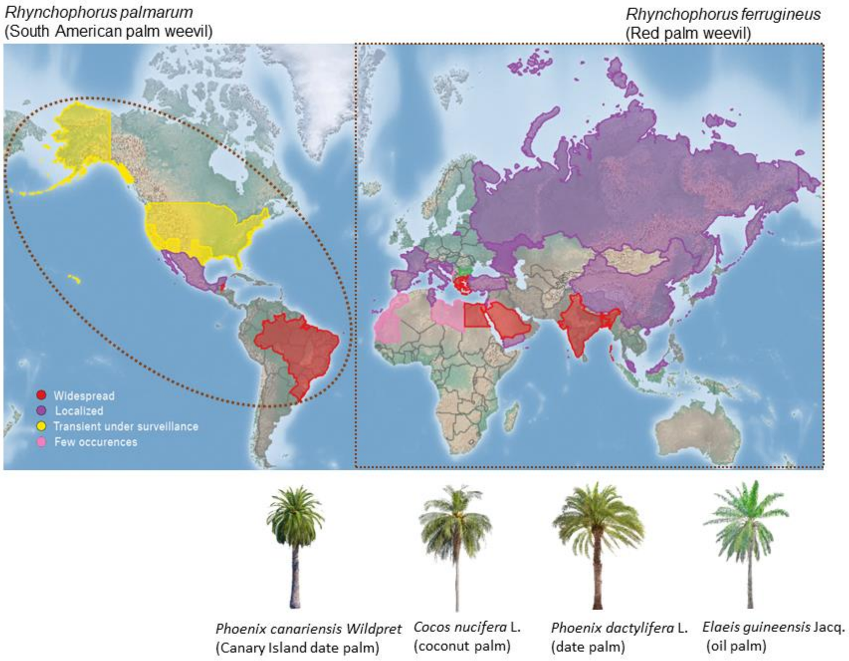

**Fig. S2** Predicted transmembrane regions of RferOR2/RpalOR2 clade genes by HMMTOP, TMHMM and Phobius.

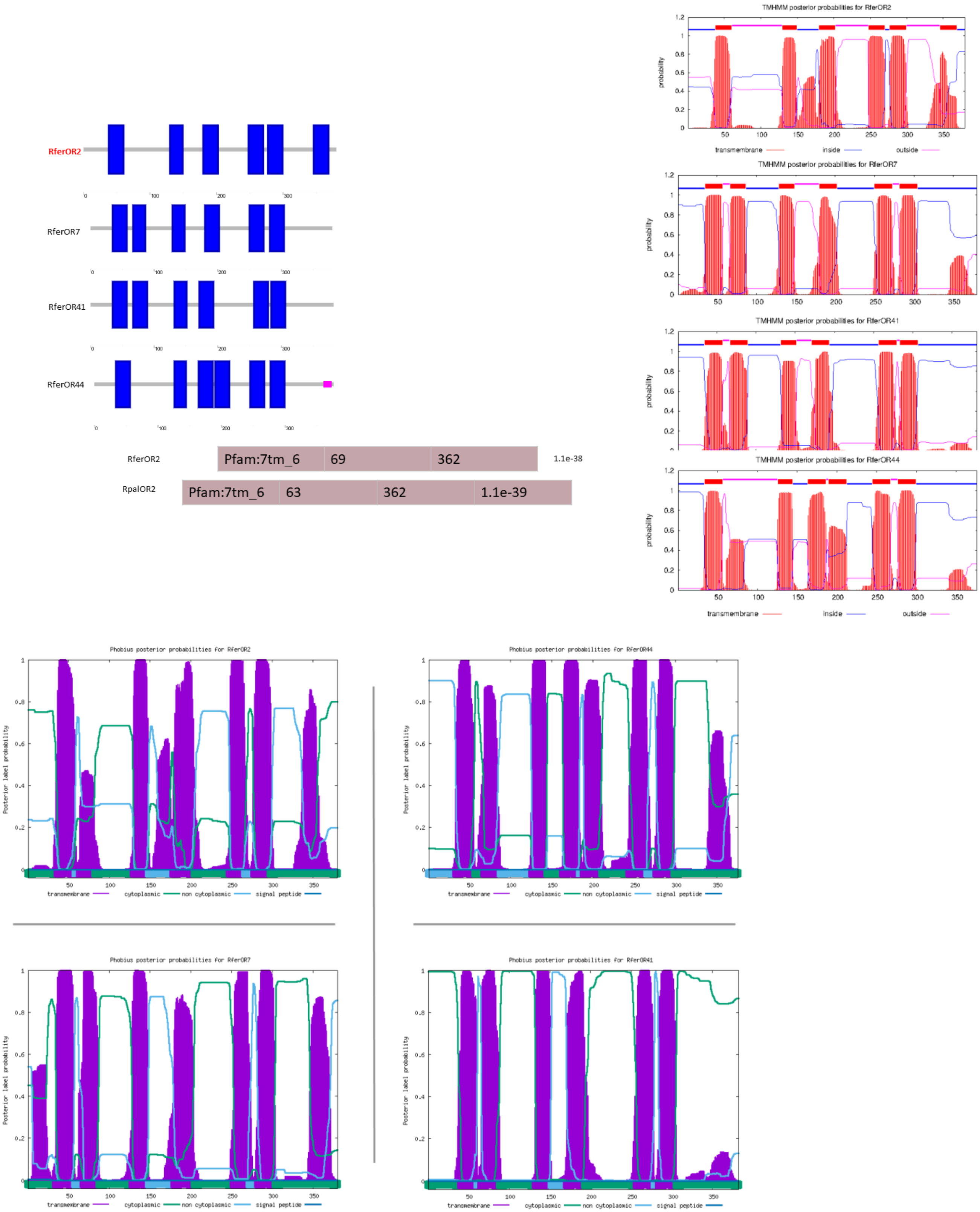

**Fig. S3 Maximum likelihood consensus tree of ORs from Coleoptera.** The tree was built from the alignment of OR amino acid sequences of *R. ferrugineus*, Rfer (red), *R. palmarum* (blue) and the following coleopteran species: *Nicrophorus vespilloides*, Nves (black); *Ips typographus*, Ityp (green); and *Megacyllene caryae*, Mcar (purple). The Orco clade was used as an outgroup. The major coleopteran OR subfamilies are indicated with blue arcs and numbers. Clades containing functionally characterized ORs pheromone and kairomone (host/plant volatile) are highlighted in yellow (*R. ferrugineus*), blue (*I. typographus* ORs), and purple (*M. caryae* ORs). Numbers on the branches are bootstrap values ($n$ = 1000). The phylogenetic tree was visualized using FigTree (http://tree.bio.ed.ac.uk/software/figtree/), and branch appearance was colored based on the bootstrap values. Scale = 3.0 amino acid substitutions per site.

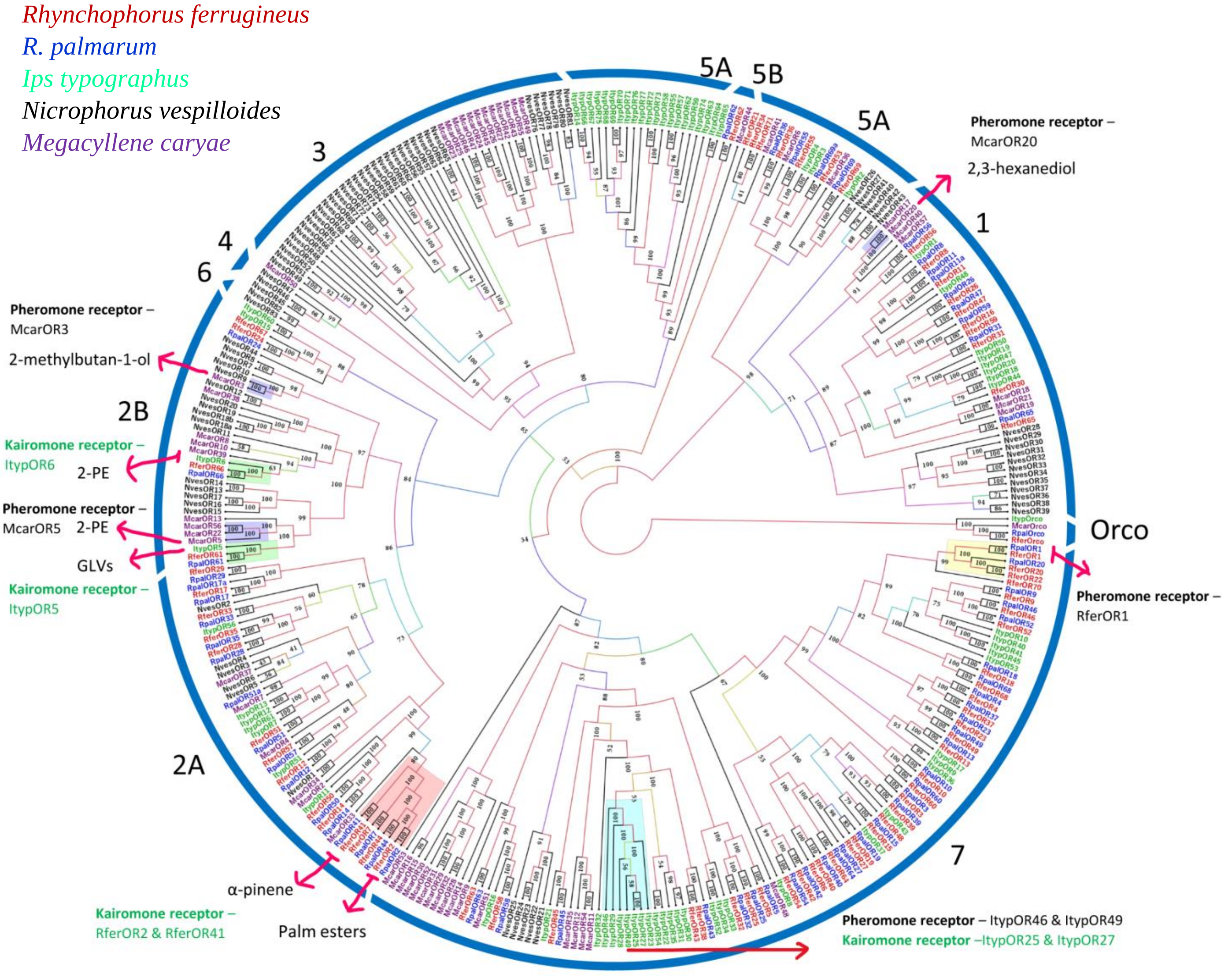

**Fig. S4** RferOR2-expressing *Drosophila* failed to respond to the RPW aggregation pheromone and structurally related compounds. Action potential frequency of *Drosophila* at1 ORNs expressing *RferOR2* when stimulated with a panel of pheromone compounds and related chemicals (100 µg loaded in the stimulus cartridge). Box plots show the number of spikes/s ±SEM.

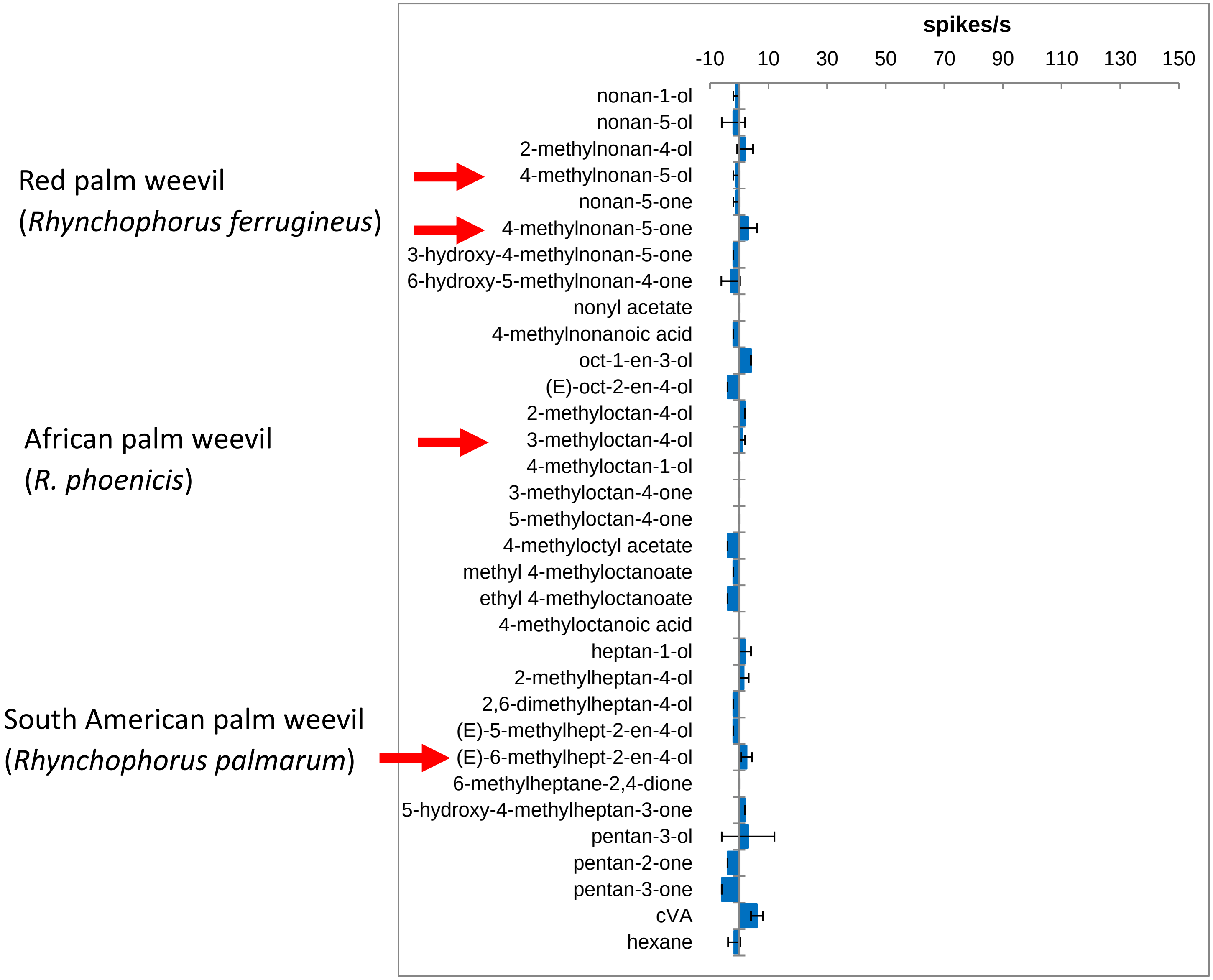

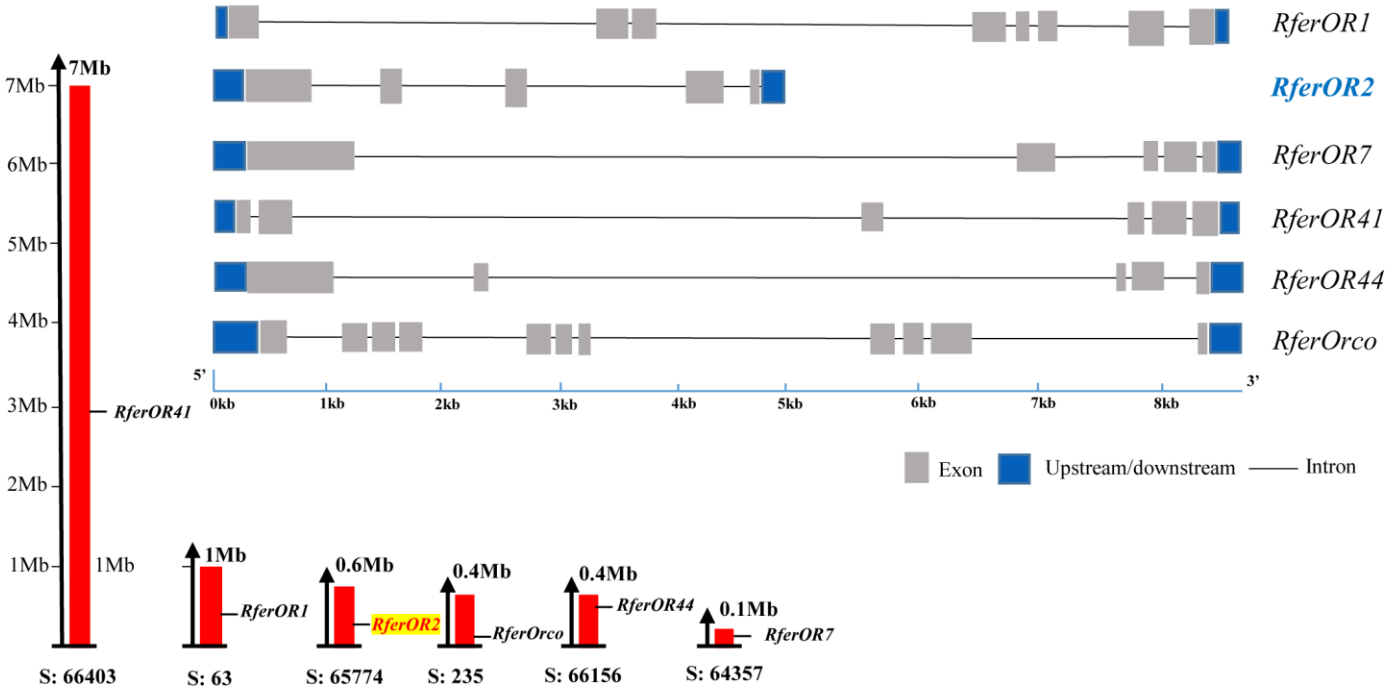


**Fig. S5** Comparison of the scaffold distribution of *R. ferrugineus* RferOR2 clade genes (in red), the pheromone receptor (RferOR1, Antony *et al*., 2021), and the odorant coreceptor (RferOrco, Soffan *et al*., 2021). Each gene's position mapped on the scaffold (S) in Mb scale was retrieved from NCBI DBSOURCE accession JAACXV010014020.1 (Dias et al., 2021). The exon-intron structure, including exon positions and gene length of RferOR2 clade genes, RferOR1, and RferOrco (upstream/downstream: 5' UTR and 3'UTR), are shown.

**Fig. S6** MEME motif analysis performed on the RferOR2 clade proteins, the pheromone receptor (RferOR1, Antony et al., 2021), and the odorant co-receptor (RferOrco, Soffan et al., 2021).

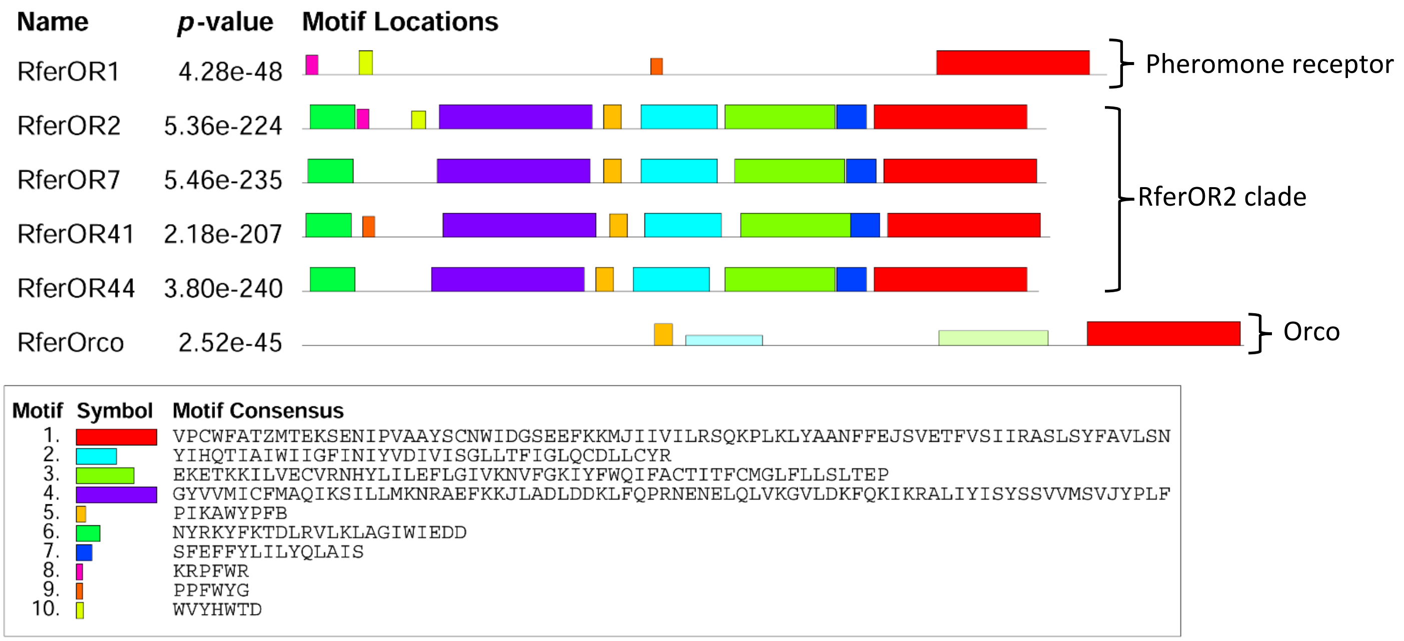

**Table S1** List of molecules tested on *Drosophila* ORNs expressing RferOR2 *via* SSR recordings.

| **Compound** | **CAS** |
|---|---|
| Hexanal | 66-25-1 |
| Nonanal | 124-19-6 |
| Isopropyl myristate | 110-27-0 |
| Propyl butyrate | 105-66-8 |
| Ethyl butyrate | 105-54-4 |
| Ethyl propionate | 105-37-3 |
| Ethyl isobutyrate | 97-62-1 |
| Ethyl acetate | 141-78-6 |
| Ethyl valerate | 539-82-2 |
| Ethyl benzoate | 93-89-0 |
| Ethyl caprylate | 106-32-1 |
| Methyl acetate | 79-20-9 |
| Methyl butyrate | 623-42-7 |
| Methyl isobutyrate | 547-63-7 |
| Butyl butyrate | 109-21-7 |
| Isobutyl propionate | 540-42-1 |
| Sec-butyl acetate/2-butyl-acetate | 105-46-4 |
| Isoamyl propionate | 105-68-0 |
| Alpha-copaene | 3856-25-5 |
| Anisole | 100-66-3 |
| Para xylene | 106-42-3 |
| Benzyl alcohol | 100-51-6 |
| 2 phenylethanol | 60-12-8 |
| Sec-butyl alcohol | 78-92-2 |
| 1 butanol | 71-36-3 |
| 3 buten-2-ol | 598-32-3 |
| 1 octen-3-ol | 3391-86-4 |
| Acetoin | 513-86-0 |
| 2 pentanone | 107-87-9 |

**Table S2.** List of aggregation pheromone compounds (with the corresponding species) and structurally related chemicals used to stimulate ORNs expressing RferOR2. All species listed are weevils belonging to the sub-family Rhynchophorinae, except *Oryctes sp*. that are palm tree feeding beetles from the family Scarabaeidae.

| **Chemicals** | **Species** | **References** |
|---|---|---|
| nonan-1-ol | - | |
| nonan-5-ol | *Metamasius hemipterus* | Ramirez *et al*. (1996) Entomol. Exp. Appl. |
| 2-methylnonan-4-ol | *Scyphophorus acupunctatus* | Ruiz-Montiel *et al.* (2003) J. Econ. Entom |
| 4-methylnonan-5-ol | *Rhynchophorus ferrugineus* | Hallett *et al*. (1993) Naturwissenschaften |
| | *Rhynchophorus bilineatus* | Oehlschlager *et al.* (1995) J. Chem. Ecol. |
| | *Rhynchophorus vulneratus* | Hallett *et al*. (1993) Naturwissenschaften |
| | *Metamasius hemipterus* | Ramirez *et al*. (1996) Entomol. Exp. Appl. |
| | *Dynamis borassi* | Giblin-Davis *et al*. (1997) J. Chem. Ecol. |
| nonan-5-one | - | |
| 4-methylnonan-5-one | *Rhynchophorus ferrugineus* | Hallett *et al*. (1993) Naturwissenschaften |
| | *Rhynchophorus vulneratus* | Hallett *et al*. (1993) Naturwissenschaften |
| 3-hydroxy-4-methylnonan-5-one | *Metamasius hemipterus* | Ramirez *et al*. (1996) Entomol. Exp. Appl. |
| 6-hydroxy-5-methylnonan-5-one | - | |
| nonyl acetate | *Oryctes elegans* | Rochat *et al*. (2004) J. Chem. Ecol. |
| 4-methylnonanoic acid | - | |
| oct-1-en-3-ol | - | |
| (*E*)-oct-2-en-4-ol | *Rhynchophorus palmarum* | Rochat *et al*. (1991) J. Chem. Ecol. |
| 2-methyloctan-4-ol | *Metamasius hemipterus* | Ramirez *et al*. (1996) Entomol. Exp. Appl. |
| | *Rhabdoscelus obscurus* | Giblin-Davis *et al*. (2000) J. Chem. Ecol. |
| | *Scyphophorus acupunctatus* | Ruiz-Montiel *et al*. (2003) J. Econ. Entom. |
| | *Sphenophorus venatus* | Duffy *et al*. (2018) J. Chem. Ecol. |
| | *Sphenophorus parvulus* | Duffy *et al*. (2018) J. Chem. Ecol. |
| | *Sphenophorus incurrens* | Illescas-Riquelme *et al*. (2016) Fla. Entomol. |
| | *Sphenophorus levis* | Zarbin *et al*. (2003) J. Chem. Ecol. |
| 4-methyloctan-1-ol | *Oryctes agamemnon* | Said *et al*. (2015) J. Chem. Ecol. |
| | *Oryctes elegans* | Rochat *et al*. (2004) J. Chem. Ecol. |
| 3-methyloctan-4-one | *Metamasius hemipterus* | Ramirez *et al.* (1996) Entomol. Exp. Appl. |
| 5-methyloctan-4-one | | |
| 4-methyloctyl acetate | *Oryctes agamemnon* | Said *et al.* (2015) J. Chem. Ecol. |
| | *Oryctes elegans* | Rochat *et al.* (2004) J. Chem. Ecol. |
| methyl 4-methyloctanoate | *Oryctes elegans* | Rochat *et al.* (2004) J. Chem. Ecol. |
| ethyl 4-methyloctanoate | *Oryctes agammemnon* | Said *et al*. (2015) J. Chem. Ecol. |
| | *Oryctes elegans* | Rochat *et al.* (2004) J. Chem. Ecol. |
| | *Oryctes monoceros* | Gries *et al*. (1994) Z. Naturforsch. C. |
| | *Oryctes rhinoceros* | Hallett *et al.* (1995) J. Chem. Ecol. |
| 4-methyloctanoic acid | *Oryctes agammemnon* | Said *et al.* (2015) J. Chem. Ecol. |
| | *Oryctes elegans* | Rochat *et al.* (2004) J. Chem. Ecol. |
| | *Oryctes monoceros* | Gries *et al.* (1994) Z. Naturforsch. C. |
| | *Oryctes rhinoceros* | Hallett *et al*. (1995) J. Chem. Ecol. |
| heptan-1-ol | - | |
| 2-methylheptan-4-ol | *Metamasius hemipterus* | Ramirez *et al*. (1996) Entomol. Exp. Appl. |
| | *Paramasius distortus* | Perez *et al*. (1997) J. Chem. Ecol. |

| | | |
|---|---|---|
| | *Rhabdoscelus obscurus* | Chang & Curtis (1972) Envion. Entomol. |
| | *Scyphophorus acupunctatus* | Ruiz-Montiel *et al*. (2003) J. Econ. Entom |
| 2,6-dimethylheptan-4-ol | - | |
| (*E*)-5-methylhept-2-en-4-ol | - | |
| (*E*)-6-methylhept-2-en-4-ol | *Rhynchophorus palmarum* | Rochat *et al*. (1991) J. Chem. Ecol. |
| | *Rhabdoscelus obscurus* | Giblin-Davis *et al*. (2000) J. Chem. Ecol. |
| 6-methylheptane-2,4-dione | - | |
| 5-hydroxy-4-methylheptan-3-one | *Sitophilus oryzae* | Phillips *et al*. (1985) J. Chem. Ecol. |
| | *Sitophilus zeamais* | Phillips *et al*. (1985) J. Chem. Ecol. |
| pentan-3-ol | *Metamasius hemipterus* | Perez *et al*. (1997) J. Chem. Ecol. |
| pentan-2-one | - | |
| pentan-3-one | *Metamasius hemipterus* | Perez *et al*. (1997) J. Chem. Ecol. |

**Table S3** List of primers used for RferOR expression studies.

| OR Primer name and Direction | Primer sequence (5' to 3') | Use |
|---|---|---|
| RferOR2_ORF_F | ATGAAGCCTGTCAAGTATCG | ORF cloning |
| RferOR2_ORF_R | CTATATAGATAACTGATTTTC | ” |
| RferOR2_PW_F1 | ACAGCCTCGGAACTCATCGG | Primer walk, sequencing |
| RferOR2_EcoR1_ORF_F | GAATTCATGAAGCCTGTCAAGTATCGTGAATTG | Dm |
| RferOR2_NotI_ORF_R | GCGGCCGCCTATATAGATAACTGATTTTCTGATGC | Dm |
| RferOR02_F | GGAAAATTTCACCACGCACCA | RACE/Dm |
| RferOR02_R | TTTTGACGTCGGCAACACTCA | RACE/Dm |
| RferOR7_walk1 | CACTGTTACTCATACCGTTGT | Primer walk, sequencing |
| RferOR7_walk2 | GCTTGACAGAACCAACAACATT | Primer walk, sequencing |
| RferOR07_F | CTGTTACTCATACCGTTGTTC | Primer walk, sequencing |
| RferOR07_R | TCGCACTGTAAGCCTATAA | Primer walk, sequencing |
| RferOR41_F | GATCACTGACGTACTTCTGCT | Primer walk, sequencing |
| RferOR41_R | GGCGAGCAATGTTATAGCGA | Primer walk, sequencing |
| RferOR47_F | TTGGCCGAAGACATTGAAGG | Primer walk, sequencing |
| RferOR47_R | TCACCGAAACTGTGGTGAAGA | Primer walk, sequencing |

*Dm: *Drosophila melanogaster*

**Table S4.** Red palm weevil antennal transcriptome assembly report.

| | Male field | Female field | Male lab | Female lab |
|---|---|---|---|---|
| Total number of raw reads | 99,146,450 | 134,645,400 | 90,215,866 | 198.956.016 |
| Total length of reads (bp) | 14,971,113,950 | 20,331,455,400 | 13,622,595,766 | 30.042.358.416 |
| Total number of cleaned reads | 98,868,937 | 134,414,781 | 90,122,367 | 198,763,141 |
| Total length of cleaned reads (bp) | 12,753,517,786 | 17,740,681,985 | 11,867,016,957 | 26,492,600,964 |
| Number of contigs | 53,645 | 59,627 | 50,519 | 81,862 |
| Total length | 31,354,749 | 35,944,586 | 30,782,692 | 45,247,906 |
| -N50 | 772 | 860 | 822 | 656 |
| Average | 584 | 603 | 609 | 553 |
| -Min | 74 | 104 | 85 | 101 |
| -Max | 21,051 | 25,370 | 14,929 | 21,384 |

**Table S4.** Red palm weevil snout transcriptome assembly report.

| | Male field | Female field | Male lab | Female lab |
|---|---|---|---|---|
| Total number of raw reads | 133,189,166 | 254,584,320 | 102,484,022 | 200,404,720 |
| Total length of reads (bp) | 20,111,564,066 | 38,442,232,320 | 15,475,087,322 | 30,261,112,720 |
| Total number of reads cleaned | 133,129,330 | 254,460,904 | 102,403,197 | 200,277,342 |
| Total length of reads cleaned (bp) | 18,172,140,980 | 34,453,144,617 | 13,685,276,150 | 26,990,967,531 |
| Number of contigs | 126,191 | 63,128 | 62,641 | 120,850 |
| Total length | 64,605,550 | 37,133,236 | 37,424,798 | 69,131,965 |
| -N50 | 588 | 792 | 777 | 702 |
| Average | 512 | 588 | 597 | 572 |
| -Min | 104 | 87 | 110 | 72 |
| -Max | 34,191 | 30,694 | 13,256 | 21,750 |

**Table S5**: Genome-wide analysis and expression profiling of antennal specific ORs (RferOR1-RferOR22) and RferOR2 clade on RPW male antennal transcriptome.

| Lab | Name (Dias et al., 2021) | Chromosome | Region (start - end) | Expression value | TPM | RPKM | Exons | Gene length | Unique gene reads | Total gene reads | Transcripts annotated | Uniquely identified transcripts | Exon length | Unique exon reads | Total exon reads | Ratio of unique to total (exon reads) | Unique exon-exon reads | Total exon-exon reads | Unique intron reads | Total intron reads | Ratio of intron to total gene reads |
|---|---|---|---|---|---|---|---|---|---|---|---|---|---|---|---|---|---|---|---|---|---|
| RferOR1 | GWI33_012310 | JAACXV010000062 | complement(297265..305118 | 102.849383 | 102.8494 | 74.40133 | 8 | 7854 | 3038 | 3038 | 1 | 1 | 1047 | 2995 | 2995 | 1 | 2564 | 2564 | 43 | 43 | 0.014154 |
| RferOR2 | GWI33_016023 | JAACXV010014020 | 299313..303900 | 27.64541737 | 27.64542 | 19.99872 | 6 | 4588 | 1105 | 1105 | 1 | 1 | 939 | 722 | 722 | 1 | 517 | 517 | 383 | 383 | 0.346606 |
| RferOR3 | GWI33_019765 | JAACXV010014478 | 352544..376822 | 5.82215466 | 5.822155 | 4.211752 | 14 | 24279 | 362 | 362 | 1 | 1 | 2112 | 342 | 342 | 1 | 245 | 245 | 20 | 20 | 0.055249 |
| RferOR4 | GWI33_020264 | JAACXV010014546 | complement(322636..324740 | 10.1839634 | 10.18396 | 7.367088 | 5 | 2105 | 331 | 331 | 1 | 1 | 1158 | 328 | 328 | 1 | 235 | 235 | 3 | 3 | 0.009063 |
| RferOR5 | GWI33_010244 | JAACXV010000048 | complement(164789..178873 | 12.66540369 | 12.6654 | 9.162164 | 8 | 14085 | 441 | 441 | 1 | 1 | 1215 | 428 | 428 | 1 | 291 | 291 | 13 | 13 | 0.029478 |
| RferOR6 | GWI33_011596 | JAACXV010009827 | 1066..7039 | 6.197936927 | 6.197937 | 4.483593 | 8 | 5974 | 278 | 278 | 1 | 1 | 1137 | 196 | 196 | 1 | 174 | 174 | 82 | 82 | 0.294964 |
| RferOR7 | GWI33_012923 | JAACXV010012833 | 16775..24411 | 2.031319695 | 2.03132 | 1.469458 | 6 | 7637 | 66 | 66 | 1 | 1 | 1062 | 60 | 60 | 1 | 41 | 41 | 6 | 6 | 0.090909 |
| RferOR8 | GWI33_011611 | JAACXV010009936 | 2902..8227 | 12.00173781 | 12.00174 | 8.682068 | 6 | 5326 | 257 | 257 | 1 | 1 | 707 | 236 | 236 | 1 | 207 | 207 | 21 | 21 | 0.081712 |
| RferOR9 | GWI33_017967 | JAACXV010014285 | 249744..253840 | 9.182373507 | 9.182374 | 6.642537 | 5 | 4097 | 172 | 172 | 1 | 1 | 603 | 154 | 154 | 1 | 124 | 124 | 18 | 18 | 0.104651 |
| RferOR10 | GWI33_017171 | JAACXV010014182 | complement(27471..36240) | 2.99619655 | 2.996197 | 2.167451 | 6 | 8770 | 69 | 69 | 1 | 1 | 780 | 65 | 65 | 1 | 57 | 57 | 4 | 4 | 0.057971 |
| RferOR11 | GWI33_019361 | JAACXV010014429 | 23897..31121 | 4.549198951 | 4.549199 | 3.290894 | 8 | 7225 | 163 | 163 | 1 | 1 | 1146 | 145 | 145 | 1 | 118 | 118 | 18 | 18 | 0.110429 |
| RferOR12 | GWI33_003915 | JAACXV010000020 | 1313556..1324879 | 1.420162527 | 1.420163 | 1.027347 | 8 | 11324 | 47 | 47 | 1 | 1 | 1038 | 41 | 41 | 1 | 35 | 35 | 6 | 6 | 0.12766 |
| RferOR13 | GWI33_021640 | JAACXV010000070 | 1302726..1316295 | 4.515044109 | 4.515044 | 3.266187 | 7 | 13570 | 162 | 163 | 2 | 1 | 1083 | 136 | 136 | 1 | 113 | 113 | 26 | 27 | 0.165644 |
| RferOR14 | GWI33_020983 | JAACXV010014602 | 6285..29953 | 0.986279164 | 0.986279 | 0.713475 | 6 | 23669 | 49 | 49 | 1 | 1 | 1203 | 33 | 33 | 1 | 15 | 15 | 16 | 16 | 0.326531 |
| RferOR15 | GWI33_017013 | JAACXV010014152 | complement(224681..228563 | 1.111990472 | 1.11199 | 0.804415 | 7 | 3883 | 39 | 39 | 1 | 1 | 1164 | 36 | 36 | 1 | 24 | 24 | 3 | 3 | 0.076923 |
| RferOR16 | GWI33_019360 | JAACXV010014429 | 5938..18339 | 1.975514209 | 1.975514 | 1.429089 | 8 | 12402 | 62 | 62 | 1 | 1 | 1092 | 60 | 60 | 1 | 48 | 48 | 2 | 2 | 0.032258 |
| RferOR17 | GWI33_005758 | JAACXV010000286 | complement(769467..772325 | 4.025424373 | 4.025424 | 2.911995 | 6 | 2859 | 94 | 94 | 1 | 1 | 786 | 88 | 88 | 1 | 74 | 74 | 6 | 6 | 0.06383 |
| RferOR18 | GWI33_013599 | JAACXV010013333 | 34829..44725 | 2.119260975 | 2.119261 | 1.533075 | 6 | 9897 | 81 | 82 | 1 | 1 | 984 | 58 | 58 | 1 | 47 | 47 | 23 | 24 | 0.292683 |
| RferOR19* | GWI33_017011 | JAACXV010014152 | complement(65172..69476) | 9.500892688 | 9.500893 | 6.872954 | 9 | 4305 | 307 | 307 | 1 | 1 | 1158 | 306 | 306 | 1 | 273 | 273 | 1 | 1 | 0.003257 |
| RferOR20 | GWI33_012311 | JAACXV010000062 | complement(312682..332296 | 1.609068518 | 1.609069 | 1.164002 | 9 | 19615 | 66 | 66 | 1 | 1 | 1296 | 58 | 58 | 1 | 51 | 51 | 8 | 8 | 0.121212 |
| RferOR21 | GWI33_004867 | JAACXV010000244 | complement(834805..839312 | 7.997452389 | 7.997452 | 5.785364 | 10 | 4508 | 396 | 396 | 1 | 1 | 1578 | 351 | 351 | 1 | 315 | 315 | 45 | 45 | 0.113636 |
| RferOR22 | GWI33_012309 | JAACXV010000062 | complement(275646..286114 | 12.36479162 | 12.36479 | 8.944701 | 8 | 10469 | 578 | 578 | 1 | 1 | 1230 | 423 | 423 | 1 | 385 | 385 | 155 | 155 | 0.268166 |
| RferOR41 | GWI33_020752 | JAACXV010014584 | complement(1001816..10099 | 4.66521879 | 4.665219 | 3.374823 | 6 | 8102 | 160 | 160 | 1 | 1 | 894 | 116 | 116 | 1 | 89 | 89 | 44 | 44 | 0.275 |
| RferOR44 | GWI33_018869 | JAACXV010014364 | complement(40098..46542) | 1.589494191 | 1.589494 | 1.149841 | 5 | 6445 | 56 | 56 | 1 | 1 | 1131 | 50 | 50 | 1 | 24 | 24 | 6 | 6 | 0.107143 |
| RferOrco | GWI33_004518 | JAACXV010000234 | 296743..304746 | 245.4263804 | 245.4264 | 177.5417 | 11 | 8004 | 9062 | 9062 | 1 | 1 | 1305 | 8908 | 8908 | 1 | 7969 | 7969 | 154 | 154 | 0.016994 |
| **Field** | | | | | | | | | | | | | | | | | | | | | |
| RferOR1 | GWI33_012310 | JAACXV010000062 | complement(297265..305118 | 41.39802258 | 41.39802 | 26.64679 | 8 | 7854 | 1130 | 1130 | 1 | 1 | 1047 | 1102 | 1102 | 1 | 931 | 931 | 28 | 28 | 0.024779 |
| RferOR2 | GWI33_016023 | JAACXV010014020 | 299313..303900 | 14.40912253 | 14.40912 | 9.274763 | 6 | 4588 | 527 | 527 | 1 | 1 | 939 | 344 | 344 | 1 | 267 | 267 | 183 | 183 | 0.347249 |
| RferOR3 | GWI33_019765 | JAACXV010014478 | 352544..376822 | 3.445263941 | 3.445264 | 2.217623 | 14 | 24279 | 197 | 197 | 1 | 1 | 2112 | 185 | 185 | 1 | 146 | 146 | 12 | 12 | 0.060914 |
| RferOR4 | GWI33_020264 | JAACXV010014546 | complement(322636..324740 | 4.517391869 | 4.517392 | 2.907723 | 5 | 2105 | 133 | 133 | 1 | 1 | 1158 | 133 | 133 | 1 | 93 | 93 | 0 | 0 | 0 |
| RferOR5 | GWI33_010244 | JAACXV010000048 | complement(164789..178873 | 6.539127055 | 6.539127 | 4.209059 | 8 | 14085 | 213 | 213 | 1 | 1 | 1215 | 202 | 202 | 1 | 134 | 134 | 11 | 11 | 0.051643 |
| RferOR6 | GWI33_011596 | JAACXV010009827 | 1066..7039 | 3.4246754 | 3.424675 | 2.204371 | 8 | 5974 | 152 | 152 | 1 | 1 | 1137 | 99 | 99 | 1 | 80 | 80 | 53 | 53 | 0.348684 |
| RferOR7 | GWI33_012923 | JAACXV010012833 | 16775..24411 | 1.70364067 | 1.703641 | 1.096587 | 6 | 7637 | 52 | 52 | 1 | 1 | 1062 | 46 | 46 | 1 | 32 | 32 | 6 | 6 | 0.115385 |
| RferOR8 | GWI33_011611 | JAACXV010009936 | 2902..8227 | 5.22941519 | 5.229415 | 3.366033 | 6 | 5326 | 112 | 112 | 1 | 1 | 707 | 94 | 94 | 1 | 81 | 81 | 18 | 18 | 0.160714 |
| RferOR9 | GWI33_017967 | JAACXV010014285 | 249744..253840 | 2.739533796 | 2.739534 | 1.763364 | 5 | 4097 | 44 | 44 | 1 | 1 | 603 | 42 | 42 | 1 | 34 | 34 | 2 | 2 | 0.045455 |
| RferOR10 | GWI33_017171 | JAACXV010014182 | complement(27471..36240) | 1.260637118 | 1.260637 | 0.811438 | 6 | 8770 | 28 | 28 | 1 | 1 | 780 | 25 | 25 | 1 | 20 | 20 | 3 | 3 | 0.107143 |
| RferOR11 | GWI33_019361 | JAACXV010014429 | 23897..31121 | 2.882964885 | 2.882965 | 1.855687 | 8 | 7225 | 105 | 105 | 1 | 1 | 1146 | 84 | 84 | 1 | 74 | 74 | 21 | 21 | 0.2 |
| RferOR12 | GWI33_003915 | JAACXV010000020 | 1313556..1324879 | 0.83362362 | 0.833624 | 0.536581 | 8 | 11324 | 25 | 25 | 1 | 1 | 1038 | 22 | 22 | 1 | 20 | 20 | 3 | 3 | 0.12 |
| RferOR13 | GWI33_021640 | JAACXV010000070 | 1302726..1316295 | 2.288003988 | 2.288004 | 1.472726 | 7 | 13570 | 74 | 75 | 2 | 1 | 1083 | 63 | 63 | 1 | 54 | 54 | 11 | 12 | 0.16 |
| RferOR14 | GWI33_020983 | JAACXV010014602 | 6285..29953 | 0.751981044 | 0.751981 | 0.48403 | 6 | 23669 | 35 | 35 | 1 | 1 | 1203 | 23 | 23 | 1 | 8 | 8 | 12 | 12 | 0.342857 |
| RferOR15 | GWI33_017013 | JAACXV010014152 | complement(224681..228563 | 0.439273552 | 0.439274 | 0.282749 | 7 | 3883 | 16 | 16 | 1 | 1 | 1164 | 13 | 13 | 1 | 7 | 7 | 3 | 3 | 0.1875 |
| RferOR16 | GWI33_019360 | JAACXV010014429 | 5938..18339 | 1.116564304 | 1.116564 | 0.718702 | 8 | 12402 | 31 | 31 | 1 | 1 | 1092 | 31 | 31 | 1 | 28 | 28 | 0 | 0 | 0 |
| RferOR17 | GWI33_005758 | JAACXV010000286 | complement(769467..772325 | 0.900730032 | 0.90073 | 0.579776 | 6 | 2859 | 19 | 19 | 1 | 1 | 786 | 18 | 18 | 1 | 11 | 11 | 1 | 1 | 0.052632 |
| RferOR18 | GWI33_013599 | JAACXV010013333 | 34829..44725 | 0.319771366 | 0.319771 | 0.205828 | 6 | 9897 | 10 | 11 | 1 | 1 | 984 | 8 | 8 | 1 | 6 | 6 | 2 | 3 | 0.272727 |
| RferOR19* | GWI33_017011 | JAACXV010014152 | complement(65172..69476) | 9.238575852 | 9.238576 | 5.946622 | 9 | 4305 | 276 | 276 | 1 | 1 | 1158 | 272 | 272 | 1 | 237 | 237 | 4 | 4 | 0.014493 |
| RferOR20 | GWI33_012311 | JAACXV010000062 | complement(312682..332296 | 0.576624756 | 0.576625 | 0.371158 | 9 | 19615 | 25 | 25 | 1 | 1 | 1296 | 19 | 19 | 1 | 17 | 17 | 6 | 6 | 0.24 |
| RferOR21 | GWI33_004867 | JAACXV010000244 | complement(834805..839312 | 2.467589309 | 2.467589 | 1.588321 | 10 | 4508 | 122 | 122 | 1 | 1 | 1578 | 99 | 99 | 1 | 83 | 83 | 23 | 23 | 0.188525 |
| RferOR22 | GWI33_012309 | JAACXV010000062 | complement(275646..286114 | 5.500067503 | 5.500068 | 3.540245 | 8 | 10469 | 203 | 203 | 1 | 1 | 1230 | 172 | 172 | 1 | 154 | 154 | 31 | 31 | 0.152709 |
| RferOR41 | GWI33_020752 | JAACXV010014584 | complement(1001816..10099 | 1.539838627 | 1.539839 | 0.991152 | 6 | 8102 | 40 | 40 | 1 | 1 | 894 | 35 | 35 | 1 | 27 | 27 | 5 | 5 | 0.125 |
| RferOR44 | GWI33_018869 | JAACXV010014364 | complement(40098..46542) | 1.008509694 | 1.00851 | 0.64915 | 5 | 6445 | 38 | 38 | 1 | 1 | 1131 | 29 | 29 | 1 | 18 | 18 | 9 | 9 | 0.236842 |
| RferOrco | GWI33_004518 | JAACXV010000234 | 296743..304746 | 122.6370972 | 122.6371 | 78.93818 | 11 | 8004 | 4155 | 4155 | 1 | 1 | 1305 | 4069 | 4069 | 1 | 3626 | 3626 | 86 | 86 | 0.020698 |

***RferOR19 and RferOR27 map in the same gene model (Dias et al., 2021).**

**Table S6**: Genome-wide analysis and expression profiling of antennal specific ORs (RferOR1-RferOR22) and RferOR2 clade on RPW female antennal transcriptome.

| Lab | Name (Dias et al., 2021) | Chromosome | Region | Expression value | TPM | RPKM | Exons | Gene length | Unique gene reads | Total gene reads | Transcripts annotated | Uniquely identified transcripts | Exon length | Unique exon reads | Total exon reads | Ratio of unique to total (exon reads) | Unique exon-exon reads | Total exon-exon reads | Unique intron reads | Total intron reads | Ratio of intron to total gene reads |
|---|---|---|---|---|---|---|---|---|---|---|---|---|---|---|---|---|---|---|---|---|---|
| RferOR1 | GWI33_012310 | JAACXV010000062 | complement(297265..305 | 89.96892073 | 89.96892 | 62.51843 | 8 | 7854 | 5436 | 5436 | 1 | 1 | 1047 | 5354 | 5354 | 1 | 4771 | 4771 | 82 | 82 | 0.015085 |
| RferOR2 | GWI33_016023 | JAACXV010014020 | 299313..303900 | 22.50288707 | 22.50289 | 15.63701 | 6 | 4588 | 2039 | 2039 | 1 | 1 | 939 | 1201 | 1201 | 1 | 878 | 878 | 838 | 838 | 0.410986 |
| RferOR3 | GWI33_019765 | JAACXV010014478 | 352544..376822 | 4.715017868 | 4.715018 | 3.276415 | 14 | 24279 | 587 | 587 | 1 | 1 | 2112 | 566 | 566 | 1 | 440 | 440 | 21 | 21 | 0.035775 |
| RferOR4 | GWI33_020264 | JAACXV010014546 | complement(322636..324 | 5.94058239 | 5.940582 | 4.128046 | 5 | 2105 | 403 | 403 | 1 | 1 | 1158 | 391 | 391 | 1 | 270 | 270 | 12 | 12 | 0.029777 |
| RferOR5 | GWI33_010244 | JAACXV010000048 | complement(164789..178 | 7.356110762 | 7.356111 | 5.111681 | 8 | 14085 | 544 | 544 | 1 | 1 | 1215 | 508 | 508 | 1 | 318 | 318 | 36 | 36 | 0.066176 |
| RferOR6 | GWI33_011596 | JAACXV010009827 | 1066..7039 | 1.887818299 | 1.887818 | 1.311824 | 8 | 5974 | 168 | 168 | 1 | 1 | 1137 | 122 | 122 | 1 | 95 | 95 | 46 | 46 | 0.27381 |
| RferOR7 | GWI33_012923 | JAACXV010012833 | 16775..24411 | 0.612968325 | 0.612968 | 0.425945 | 6 | 7637 | 45 | 45 | 1 | 1 | 1062 | 37 | 37 | 1 | 26 | 26 | 8 | 8 | 0.177778 |
| RferOR8 | GWI33_011611 | JAACXV010009936 | 2902..8227 | 5.872910935 | 5.872911 | 4.081022 | 6 | 5326 | 258 | 258 | 1 | 1 | 707 | 236 | 236 | 1 | 207 | 207 | 22 | 22 | 0.085271 |
| RferOR9 | GWI33_017967 | JAACXV010014285 | 249744..253840 | 6.156387797 | 6.156388 | 4.278007 | 5 | 4097 | 242 | 242 | 1 | 1 | 603 | 211 | 211 | 1 | 167 | 167 | 31 | 31 | 0.128099 |
| RferOR10 | GWI33_017171 | JAACXV010014182 | complement(27471..3624 | 1.037585884 | 1.037586 | 0.721007 | 6 | 8770 | 73 | 73 | 1 | 1 | 780 | 46 | 46 | 1 | 42 | 42 | 27 | 27 | 0.369863 |
| RferOR11 | GWI33_019361 | JAACXV010014429 | 23897..31121 | 2.778783956 | 2.778784 | 1.930947 | 8 | 7225 | 224 | 224 | 1 | 1 | 1146 | 181 | 181 | 1 | 144 | 144 | 43 | 43 | 0.191964 |
| RferOR12 | GWI33_003915 | JAACXV010000020 | 1313556..1324879 | 0.593241489 | 0.593241 | 0.412237 | 8 | 11324 | 44 | 44 | 1 | 1 | 1038 | 35 | 35 | 1 | 32 | 32 | 9 | 9 | 0.204545 |
| RferOR13 | GWI33_021640 | JAACXV010000070 | 1302726..1316295 | 1.884474904 | 1.884475 | 1.309501 | 7 | 13570 | 134 | 135 | 2 | 1 | 1083 | 116 | 116 | 1 | 102 | 102 | 18 | 19 | 0.140741 |
| RferOR14 | GWI33_020983 | JAACXV010014602 | 6285..29953 | 0.921373565 | 0.921374 | 0.640252 | 6 | 23669 | 108 | 108 | 1 | 1 | 1203 | 63 | 63 | 1 | 30 | 30 | 45 | 45 | 0.416667 |
| RferOR15 | GWI33_017013 | JAACXV010014152 | complement(224681..228 | 0.151149893 | 0.15115 | 0.105032 | 7 | 3883 | 17 | 17 | 1 | 1 | 1164 | 10 | 10 | 1 | 7 | 7 | 7 | 7 | 0.411765 |
| RferOR16 | GWI33_019360 | JAACXV010014429 | 5938..18339 | 1.450042384 | 1.450042 | 1.007619 | 8 | 12402 | 93 | 93 | 1 | 1 | 1092 | 90 | 90 | 1 | 75 | 75 | 3 | 3 | 0.032258 |
| RferOR17 | GWI33_005758 | JAACXV010000286 | complement(769467..772 | 2.238403001 | 2.238403 | 1.555442 | 6 | 2859 | 108 | 108 | 1 | 1 | 786 | 100 | 100 | 1 | 92 | 92 | 8 | 8 | 0.074074 |
| RferOR18 | GWI33_013599 | JAACXV010013333 | 34829..44725 | 0.572157645 | 0.572158 | 0.397586 | 6 | 9897 | 53 | 53 | 1 | 1 | 984 | 32 | 32 | 1 | 21 | 21 | 21 | 21 | 0.396226 |
| RferOR19* | GWI33_017011 | JAACXV010014152 | complement(65172..6947 | 5.91019578 | 5.910196 | 4.106931 | 9 | 4305 | 392 | 392 | 1 | 1 | 1158 | 389 | 389 | 1 | 359 | 359 | 3 | 3 | 0.007653 |
| RferOR20 | GWI33_012311 | JAACXV010000062 | complement(312682..332 | 0.570170987 | 0.570171 | 0.396206 | 9 | 19615 | 56 | 56 | 1 | 1 | 1296 | 42 | 42 | 1 | 33 | 33 | 14 | 14 | 0.25 |
| RferOR21 | GWI33_004867 | JAACXV010000244 | complement(834805..839 | 7.31404564 | 7.314046 | 5.082451 | 10 | 4508 | 737 | 737 | 1 | 1 | 1578 | 656 | 656 | 1 | 600 | 600 | 81 | 81 | 0.109905 |
| RferOR22 | GWI33_012309 | JAACXV010000062 | complement(275646..286 | 7.180578448 | 7.180578 | 4.989706 | 8 | 10469 | 711 | 711 | 1 | 1 | 1230 | 502 | 502 | 1 | 482 | 482 | 209 | 209 | 0.293952 |
| RferOR41 | GWI33_020752 | JAACXV010014584 | complement(1001816..10 | 1.515353763 | 1.515354 | 1.053003 | 6 | 8102 | 107 | 107 | 1 | 1 | 894 | 77 | 77 | 1 | 65 | 65 | 30 | 30 | 0.280374 |
| RferOR44 | GWI33_018869 | JAACXV010014364 | complement(40098..4654 | 0.777800512 | 0.777801 | 0.540485 | 5 | 6445 | 54 | 54 | 1 | 1 | 1131 | 50 | 50 | 1 | 35 | 35 | 4 | 4 | 0.074074 |
| RferOrco | GWI33_004518 | JAACXV010000234 | 296743..304746 | 170.6131351 | 170.6131 | 118.5572 | 11 | 8004 | 12945 | 12945 | 1 | 1 | 1305 | 12655 | 12655 | 1 | 11405 | 11405 | 290 | 290 | 0.022402 |
| **Field** | | | | | | | | | | | | | | | | | | | | | |
| RferOR1 | GWI33_012310 | JAACXV010000062 | complement(297265..305 | 54.57584438 | 54.57584 | 36.41915 | 8 | 7854 | 2094 | 2094 | 1 | 1 | 1047 | 2055 | 2055 | 1 | 1792 | 1792 | 39 | 39 | 0.018625 |
| RferOR2 | GWI33_016023 | JAACXV010014020 | 299313..303900 | 16.5531837 | 16.55318 | 11.04615 | 6 | 4588 | 900 | 900 | 1 | 1 | 939 | 559 | 559 | 1 | 394 | 394 | 341 | 341 | 0.378889 |
| RferOR3 | GWI33_019765 | JAACXV010014478 | 352544..376822 | 3.199246319 | 3.199246 | 2.134898 | 14 | 24279 | 254 | 254 | 1 | 1 | 2112 | 243 | 243 | 1 | 187 | 187 | 11 | 11 | 0.043307 |
| RferOR4 | GWI33_020264 | JAACXV010014546 | complement(322636..324 | 4.009989458 | 4.009989 | 2.675917 | 5 | 2105 | 171 | 171 | 1 | 1 | 1158 | 167 | 167 | 1 | 99 | 99 | 4 | 4 | 0.023392 |
| RferOR5 | GWI33_010244 | JAACXV010000048 | complement(164789..178 | 6.362149017 | 6.362149 | 4.245543 | 8 | 14085 | 287 | 287 | 1 | 1 | 1215 | 278 | 278 | 1 | 198 | 198 | 9 | 9 | 0.031359 |
| RferOR6 | GWI33_011596 | JAACXV010009827 | 1066..7039 | 2.959103971 | 2.959104 | 1.974648 | 8 | 5974 | 169 | 169 | 1 | 1 | 1137 | 121 | 121 | 1 | 103 | 103 | 48 | 48 | 0.284024 |
| RferOR7 | GWI33_012923 | JAACXV010012833 | 16775..24411 | 1.38767151 | 1.387672 | 0.926011 | 6 | 7637 | 65 | 65 | 1 | 1 | 1062 | 53 | 53 | 1 | 31 | 31 | 12 | 12 | 0.184615 |
| RferOR8 | GWI33_011611 | JAACXV010009936 | 2902..8227 | 3.736280821 | 3.736281 | 2.493267 | 6 | 5326 | 100 | 100 | 1 | 1 | 707 | 95 | 95 | 1 | 85 | 85 | 5 | 5 | 0.05 |
| RferOR9 | GWI33_017967 | JAACXV010014285 | 249744..253840 | 3.642881954 | 3.642882 | 2.430941 | 5 | 4097 | 88 | 88 | 1 | 1 | 603 | 79 | 79 | 1 | 63 | 63 | 9 | 9 | 0.102273 |
| RferOR10 | GWI33_017171 | JAACXV010014182 | complement(27471..3624 | 0.819914473 | 0.819914 | 0.547139 | 6 | 8770 | 27 | 27 | 1 | 1 | 780 | 23 | 23 | 1 | 19 | 19 | 4 | 4 | 0.148148 |
| RferOR11 | GWI33_019361 | JAACXV010014429 | 23897..31121 | 1.334484061 | 1.334484 | 0.890518 | 8 | 7225 | 67 | 67 | 1 | 1 | 1146 | 55 | 55 | 1 | 49 | 49 | 12 | 12 | 0.179104 |
| RferOR12 | GWI33_003915 | JAACXV010000020 | 1313556..1324879 | 0.642908559 | 0.642909 | 0.429021 | 8 | 11324 | 33 | 33 | 1 | 1 | 1038 | 24 | 24 | 1 | 19 | 19 | 9 | 9 | 0.272727 |
| RferOR13 | GWI33_021640 | JAACXV010000070 | 1302726..1316295 | 1.951283871 | 1.951284 | 1.302116 | 7 | 13570 | 87 | 87 | 2 | 1 | 1083 | 76 | 76 | 1 | 52 | 52 | 11 | 11 | 0.126437 |
| RferOR14 | GWI33_020983 | JAACXV010014602 | 6285..29953 | 0.462274234 | 0.462274 | 0.308481 | 6 | 23669 | 33 | 33 | 1 | 1 | 1203 | 20 | 20 | 1 | 7 | 7 | 13 | 13 | 0.393939 |
| RferOR15 | GWI33_017013 | JAACXV010014152 | complement(224681..228 | 0.477762804 | 0.477763 | 0.318817 | 7 | 3883 | 20 | 20 | 1 | 1 | 1164 | 20 | 20 | 1 | 16 | 16 | 0 | 0 | 0 |
| RferOR16 | GWI33_019360 | JAACXV010014429 | 5938..18339 | 1.06945366 | 1.069454 | 0.71366 | 8 | 12402 | 42 | 42 | 1 | 1 | 1092 | 42 | 42 | 1 | 37 | 37 | 0 | 0 | 0 |
| RferOR17 | GWI33_005758 | JAACXV010000286 | complement(769467..772 | 0.884408243 | 0.884408 | 0.590177 | 6 | 2859 | 26 | 26 | 1 | 1 | 786 | 25 | 25 | 1 | 22 | 22 | 1 | 1 | 0.038462 |
| RferOR18 | GWI33_013599 | JAACXV010013333 | 34829..44725 | 0.791221814 | 0.791222 | 0.527992 | 6 | 9897 | 34 | 34 | 1 | 1 | 984 | 28 | 28 | 1 | 19 | 19 | 6 | 6 | 0.176471 |
| RferOR19* | GWI33_017011 | JAACXV010014152 | complement(65172..6947 | 8.260098045 | 8.260098 | 5.512068 | 9 | 4305 | 352 | 352 | 1 | 1 | 1158 | 344 | 344 | 1 | 293 | 293 | 8 | 8 | 0.022727 |
| RferOR20 | GWI33_012311 | JAACXV010000062 | complement(312682..332 | 1.072754443 | 1.072754 | 0.715863 | 9 | 19615 | 55 | 55 | 1 | 1 | 1296 | 50 | 50 | 1 | 43 | 43 | 5 | 5 | 0.090909 |
| RferOR21 | GWI33_004867 | JAACXV010000244 | complement(834805..839 | 3.101280069 | 3.10128 | 2.069523 | 10 | 4508 | 229 | 229 | 1 | 1 | 1578 | 176 | 176 | 1 | 114 | 114 | 53 | 53 | 0.231441 |
| RferOR22 | GWI33_012309 | JAACXV010000062 | complement(275646..286 | 3.75265203 | 3.752652 | 2.504192 | 8 | 10469 | 205 | 205 | 1 | 1 | 1230 | 166 | 166 | 1 | 153 | 153 | 39 | 39 | 0.190244 |
| RferOR41 | GWI33_020752 | JAACXV010014584 | complement(1001816..10 | 1.150799129 | 1.150799 | 0.767943 | 6 | 8102 | 43 | 43 | 1 | 1 | 894 | 37 | 37 | 1 | 28 | 28 | 6 | 6 | 0.139535 |
| RferOR44 | GWI33_018869 | JAACXV010014364 | complement(40098..4654 | 0.688383965 | 0.688384 | 0.459367 | 5 | 6445 | 37 | 37 | 1 | 1 | 1131 | 28 | 28 | 1 | 22 | 22 | 9 | 9 | 0.243243 |
| RferOrco | GWI33_004518 | JAACXV010000234 | 296743..304746 | 137.8570841 | 137.8571 | 91.99378 | 11 | 8004 | 6551 | 6551 | 1 | 1 | 1305 | 6470 | 6470 | 1 | 5866 | 5866 | 81 | 81 | 0.012365 |

***RferOR19 and RferOR27 map in the same gene model (Dias et al., 2021).**

**Table S7**: Expression level distribution of RferORs from the RferOR2 clade, RferOR1, and RferOrco, obtained from the transcript quantification of male and female - field vs. lab RPWs.

| Name | Chromosome | Region | Max group mean | $Log_2$ fold change | Fold change | P-value | FDR p-value | Bonferroni | Gene |
|---|---|---|---|---|---|---|---|---|---|
| GWI33_012310 | JAACXV010000062 | complement(297265..305118) | 68.45987916 | -0.609280222 | -1.52549793 | 0.153331791 | 0.531622944 | 1 | RferOR1 |
| GWI33_016023 | JAACXV010014020 | 299313..303900 | 17.81786531 | -0.291714888 | -1.224094457 | 0.468182845 | 0.830528516 | 1 | RferOR2 |
| GWI33_012923 | JAACXV010012833 | 16775..24411 | 1.011299094 | 0.710448122 | 1.636312301 | 0.394722546 | 0.785941023 | 1 | RferOR7 |
| GWI33_020752 | JAACXV010014584 | complement(1001816..1009917) | 2.213913097 | -0.724583137 | -1.652423103 | 0.373446065 | 0.770394547 | 1 | RferOR41 |
| GWI33_018869 | JAACXV010014364 | complement(40098..46542) | 0.845163302 | -0.020146966 | -1.014062776 | 0.979532108 | 0.995311429 | 1 | RferOR44 |
| GWI33_004518 | JAACXV010000234 | 296743..304746 | 148.0494352 | -0.257569763 | -1.195463235 | 0.460252845 | 0.825289623 | 1 | RferOrco |

**Table S8**: Mapping the expression level distribution of RferORs from the RferOR2 clade, RferOR1, and RferOrco, obtained from the transcript quantification of field and lab RPWs - Male vs. Femal

| Name | Chromosome | Region | Max group mean | $Log_2$ fold change | Fold change | P-value | FDR p-value | Bonferroni | Gene |
|---|---|---|---|---|---|---|---|---|---|
| GWI33_012310 | JAACXV010000062 | complement(297265..305118) | 50.52406055 | 0.032754614 | 1.022963461 | 0.967662432 | 0.99999045 | 1 | RferOR1 |
| GWI33_016023 | JAACXV010014020 | 299313..303900 | 14.6367408 | 0.177025844 | 1.130550819 | 0.799643887 | 0.99999045 | 1 | RferOR2 |
| GWI33_012923 | JAACXV010012833 | 16775..24411 | 1.283022938 | 1.029736413 | 2.041651199 | 0.402658563 | 0.99999045 | 1 | RferOR7 |
| GWI33_020752 | JAACXV010014584 | complement(1001816..1009917) | 2.182987913 | 1.238062784 | 2.358815834 | 0.280113 | 0.99999045 | 1 | RferOR41 |
| GWI33_018869 | JAACXV010014364 | complement(40098..46542) | 0.899495953 | 0.910609505 | 1.879839519 | 0.40949406 | 0.99999045 | 1 | RferOR44 |
| GWI33_004518 | JAACXV010000234 | 296743..304746 | 128.2399206 | 0.313783193 | 1.242962863 | 0.607900115 | 0.99999045 | 1 | RferOrco |

**Table S9** Identification and characteristics of *R. ferrugineus* RferOR2 clade genes in comparison to the pheromone receptor (RferOR1).

| ***R. ferrugineus* gene** | **Scaffold** (Dias et al., 2021) | ***R. palmarum* ortholog** | **ORF length (bp)** | **AA length** | **MW (kD)** | **PI** | **TM number** | **Exons** | **Ligand** |
|---|---|---|---|---|---|---|---|---|---|
| RferOR1 | 63 | RpalOR1 | 1236 | 411 | 47.47 | 6.21 | 6 | 8 | 4-methyl-5-nonanol<br>4-methyl nonanone |
| RferOR2 | 65774 | RpalOR2 | 1143 | 380 | 44.71 | 6.70 | 8 | 5 | Ethyl propionate<br>Ethyl isobutyrate<br>Ethyl acetate |
| RferOR7 | 64357 | RpalOR7 | 1143 | 380 | 44.52 | 6.93 | 8 | 5 | - |
| RferOR41 | 66403 | RpalOR41 | 1149 | 382 | 44.97 | 8.85 | 6 | 6 | α-pinene |
| RferOR44 | 66156 | RpalOR44 | 1131 | 376 | 43.68 | 6.52 | 8 | 5 | - |

-not known; ORF: open reading frame; AA: amino acids; PI: isoelectric point; TM: transmembrane.

**Table S10** Pairwise amino acid identity generated based on MAFFT multiple alignment at SIAS (http://imed.med.ucm.es/Tools/sias.html).

| | **Percentage amino acid identity** | | | | | | | |
|---|---|---|---|---|---|---|---|---|
| *RferOR2* | 100 | | | | | | | |
| *RpalOR2* | 86.76 | 100 | | | | | | |
| *RferOR44* | 38.29 | 38.82 | 100 | | | | | |
| *RpalOR44* | 35.37 | 37.05 | 78.98 | 100 | | | | |
| *RferOR7* | 38.15 | 40.00 | 45.21 | 43.35 | 100 | | | |
| *RpalOR7* | 39.21 | 39.70 | 43.35 | 42.81 | 78.94 | 100 | | |
| *RferOR41* | 30.52 | 33.23 | 37.50 | 37.50 | 35.60 | 38.68 | 100 | |
| *RpalOR41* | 31.17 | 30.90 | 50.00 | 48.48 | 49.24 | 53.78 | 85.60 | 100 |
| | *RferOR2* | *RpalOR2* | *RferOR44* | *RpalOR44* | *RferOR7* | *RpalOR7* | *RferOR41* | *RpalOR41* |

**Table S11.** List of the amino acid residues of RferOR2 active site.

| **Amino acid** | **Nature of the amino acid** |
|---|---|
| W57 | Hydrophobic |
| K65 | Positive |
| M66 | Hydrophobic |
| A68 | Hydrophobic |
| S69 | Hydrophilic |
| M70 | Hydrophobic |
| Y72 | Hydrophobic |
| M141 | Hydrophobic |
| I142 | Hydrophobic |
| Y145 | Hydrophobic |
| P146 | Hydrophobic |
| E151 | Negative |
| H152 | Positive |
| L153 | Hydrophobic |
| T154 | Hydrophilic |
| P155 | Hydrophobic |
| I156 | Hydrophobic |
| R157 | Positive |
| S158 | Hydrophilic |
| Y160 | Hydrophobic |
| N163 | Hydrophilic |
| I164 | Hydrophobic |
| N165 | Hydrophilic |
| F170 | Hydrophobic |
| F173 | Hydrophobic |
| Y265 | Hydrophobic |
| L266 | Hydrophobic |
| V269 | Hydrophobic |
| A270 | Hydrophobic |
| K273 | Positive |
| F275 | Hydrophobic |
| E276 | Negative |
| L278 | Hydrophobic |
| Y279 | Hydrophobic |

**Table S12** List of coleopteran ORs functionally characterized and detecting host/plant volatiles.

| **Coleopteran insects** | **OR** | **Host/plant volatiles** | **Host** | **Reference** |
|---|---|---|---|---|
| Eurasian spruce bark beetle,<br>*Ips typographus* L. (Ityp) | ItypOR6<br>ItypOR5<br>ItypOR27<br>ItypOR25<br>ItypOR23<br>ItypOR29 | 2-phenylethanol (2-PE)<br>Angiosperm green leaf volatiles (GLVs)<br>p-cymene<br>(+)-3-carene<br>(+)-trans-(1R, 4S)-4-thujanol<br>(+)-isopinocamphone | Spruce tree<br><br><br><br>beetle-associated fungi | Roberts et al., 2021 (7)<br><br>Hou et al., 2021(8)<br><br>Hou et al., 2021 (8) |
| The mountain pine beetle,<br>*Dendroctonus ponderosae* Hopkins (Dpon) | DponOR8<br>DponOR9 | 2-phenylethanol (2-PE),<br>Angiosperm green leaf volatiles (GLVs) | Pine trees | Roberts et al., 2021 (7) |
| The pine weevil,<br>*Hylobius abietis* (Habi) | HabiOR3<br>HabiOR4 | 2-phenylethanol (2-PE),<br>Angiosperm green leaf volatiles (GLVs) | Conifer plantations | Roberts et al., 2021 (7) |
| The red palm weevil,<br>*Rhynchophorus ferrugineus* (Rfer) | RferOR6<br>(RferOR41)<br>RferOR2 | α-pinene<br><br>Palm esters<br>2-pentanone<br>Acetoin | Palm trees | Ji et al., 2021 (9)<br><br>Current study |
| Dark Black Chafer,<br>*Holotrichia parallela* (Hpar) | HparOR27 | hexanal, lauric acid, and tetradecane | Soil pest that affects many plants | Wang et al., 2020 (10) |
| The ladybird beetle,<br>*Hippodamia variegata* (Goeze) (Hvar) | HvarOR25 | 3-hexenyl acetate, hexyl butyrate, hexyl hexanoate, benzyl alcohol, 3-methyl-3-buten-1-ol, and trans-2-hexenoic acid | coccinellid predator in cotton fields infested by *Aphis gossypii* (Glover) | Xie et al., 2022 (11) |
